# WNT-mediating TCF/LEF transcription factor gene expression in early human pluripotency and cell lineages differs from the rodent paradigm

**DOI:** 10.1101/2024.10.08.617210

**Authors:** Connor Ross, Takuya Azami, Marika Salonna, Richard Gyuris, Jennifer Nichols, Stefan Hoppler

## Abstract

Embryonic stem cell research has uncovered different requirements for WNT/β-catenin signalling in human naïve pluripotent cells compared to the mouse paradigm. It is therefore important to study WNT/β-catenin signalling directly in models of early human development. Since TCF/LEF factors mediate the regulation of target genes downstream of WNT/β-catenin signalling, we studied the expression and protein localisation of the four TCF/LEF genes by analysing *in vitro* “snapshots” of human development, leveraging naïve and primed pluripotent cells as well as extraembryonic and early embryonic cell lineages. Strikingly, we comprehensively confirm clear differences between mouse and human pluripotent stem cells, suggesting species-specific requirements for WNT signalling that may reflect differences in states of pluripotency. Human naïve ES cells express very low TCF7L1, unlike their mouse counterparts. TCF7L2 is robustly expressed in human naïve ES-derived trophectoderm cells. In human primed pluripotent stem cells, activation of WNT/β-Catenin signalling is required to induce expression of both *TCF7* and *LEF1*, concomitant with hallmark gastrulation markers. This expression of human TCF/LEF genes benchmarks differential requirements for WNT/β-catenin signalling throughout early human embryo development that requires further investigation.

## Introduction

Stem cells isolated from the early embryo provide accessible experimental models for studying early embryonic development in cell culture, especially in the context of early human development. The stable stem cell cultures originally isolated from mouse embryos (Evans and Kaufman, 1981; Martin, 1981) differ from the first human embryonic stem cells (Thomson et al., 1998) in several respects, subsequently recognised to represent different states of pluripotency; naïve and primed, respectively (Nichols and Smith, 2009). Human embryonic stem cells can be cultured in conditions supportive of naïve pluripotency (Bayerl et al., 2021; Bredenkamp et al., 2019; Guo et al., 2017; Guo et al., 2016; Khan et al., 2021; Takashima et al., 2014) but they differ from mouse naïve embryonic stem cells in their less restricted potential to differentiate into extraembryonic lineages (Guo et al., 2021; Linneberg-Agerholm et al., 2019). Some of these differences in the naïve state between human and mouse models involves WNT/β-catenin signalling. For instance, WNT activation (as part of 2i+LIF) is required in culture conditions to sustain mouse naïve embryonic stem cells (Wray et al., 2011; Ying et al., 2008). Additionally, there are differences in expression of a downstream component of the Wnt/β-catenin pathway (i.e., TCF7L1 expression, Rostovskaya et al., 2019, and see below); and, probably related to that, of Wnt/β-catenin target genes such as *Esrrb* (Martello et al., 2012).

WNT signalling is a widely conserved biochemical cell-to-cell signalling network, which mediates diverse biological functions, including during early embryonic development and later tissue stem cell-mediated regeneration and disease (reviewed by Hoppler and Moon, 2014; Nusse and Clevers, 2017). The so-called canonical or WNT/β-catenin-dependent pathway is the best understood signal transduction pathway (reviewed by Hoppler and Nakamura, 2014). WNT signal transduction in WNT receiving cells consequently drives β-Catenin protein into the nucleus to regulate the transcription of context-specific WNT target genes in association with sequence-specific DNA-binding factors, primarily of the T Cell Factor (TCF) and Lymphoid Enhancer Factor (LEF) family (reviewed by Cadigan and Waterman, 2012; Hoppler and Waterman, 2014). These transcription factors function in a bimodal fashion to either repress target gene expression without and activate them with nuclear β-Catenin (reviewed by Ramakrishnan and Cadigan, 2017). Differences between human and mouse naïve embryonic stem cells may be attributable to divergent nuclear WNT/β-catenin pathway mechanisms and particularly to differences in TCF/LEF expression and function during early embryonic development. In vertebrates, there are four TCF/LEF genes; TCF7, TCF7L1 (formerly TCF3), TCF7L2 (formerly TCF4) and LEF1 (Torres-Aguila et al., 2022), which mediate different biological functions in a cell- and context-dependent manner that are not yet understood (reviewed by Hoppler and Waterman, 2014).

To advance our understanding of the WNT signalling in early human development and how it might deviate from the mouse model-derived paradigm, we studied the expression and localisation of all four TCF/LEF transcription factors during directed differentiation from naïve and primed state pluripotent stem cells into extra-embryonic and early embryonic cell lineages, respectively. These findings confirm clear differences with mouse and provide a solid conceptional platform for future investigation into the role of WNT signalling in early human embryogenesis.

## Results

### Comparison of TCF/LEF expression between mouse and human in naïve and primed pluripotency

The mutually exclusive expression of naïve and primed markers can be used to characterise different pluripotent states. Mouse naïve embryonic stem cells robustly express Klf4 and Tcfp2l1, which is lost when cultured in primed conditions (AFX: **<u>A</u>**CTIVIN A, **<u>F</u>**GF2 and **<u>X</u>**AV9393), but both populations maintain homogeneous Oct4 expression (Fig. 1A). Human pluripotent cells in both naïve and primed states express NANOG and OCT4, whereas KLF17 is restricted to the naïve state (Fig.1B)(Blakeley et al., 2015; Stirparo et al., 2018).

**Fig. 1:**
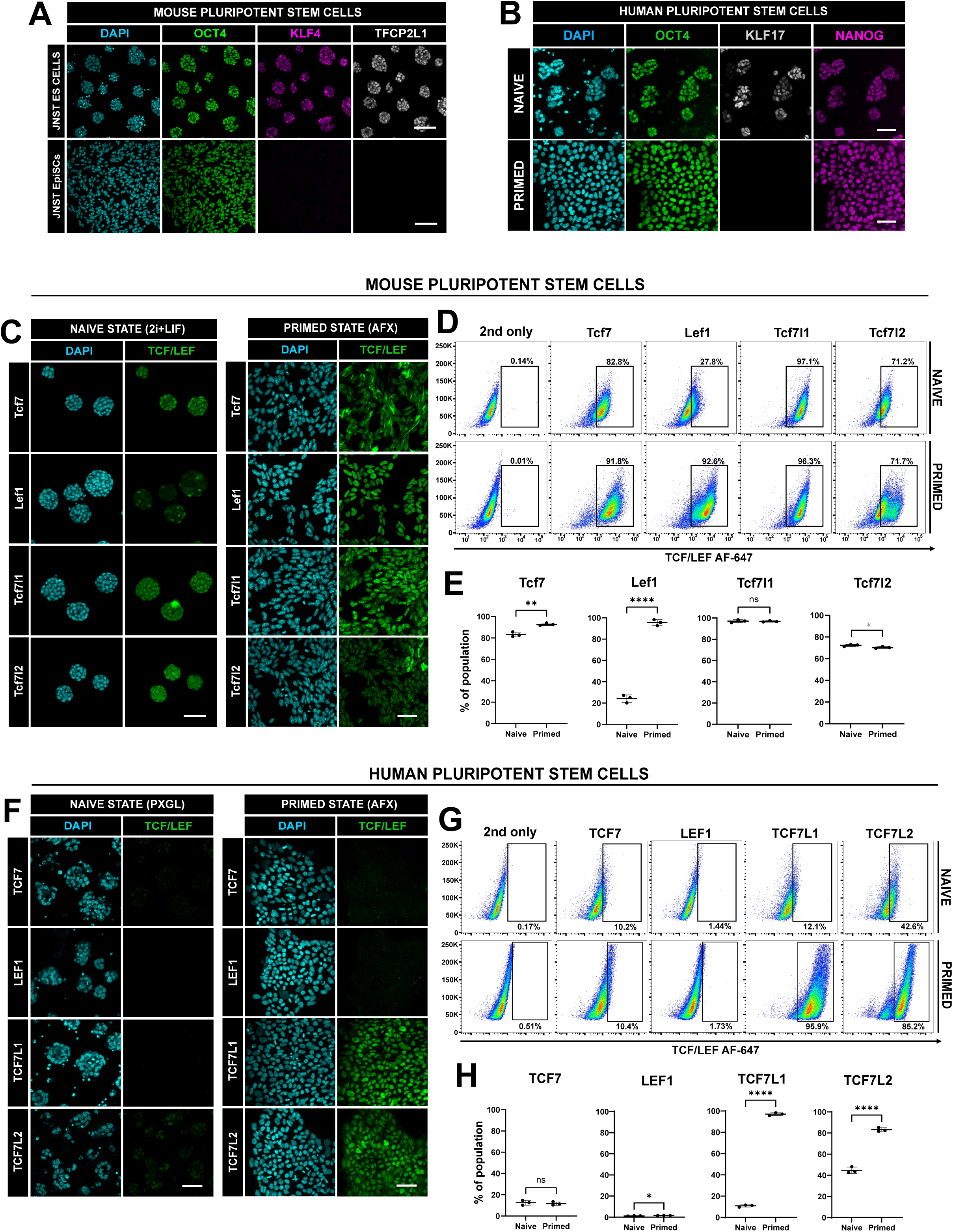
Difference in TCF/LEF expression between human and mouse in both naïve and primed embryonic stem cells. **(A)** Immunostaining of JNST mouse embryonic stem cells cultured in either 2i+LIF (naïve, top) and AFX (primed, bottom) for 48 hours for OCT4 (green), KLF4 (magenta) and TFCP2L1 (grey) with DAPI (blue). Scale bars = 100 μm. **(B)** Immunostaining of HNES1 embryonic stem cells for OCT4 (green), KLF17 (grey) and NANOG (magenta) after 72 hours of culture in PXGL (naïve) or AFX/StemFlex medium (primed). Scale bars = 100 μm. **(C)** Immunostaining of JNST mouse ES cells (2i+LIF) and HNES1 (PXGL) cells after 72 hours of culture for TCF7, TCF7L1, TCF7L2 and LEF1. Scale bars = 100 μm. **(D)** Flow cytometry of mouse naïve and primed cells for each TCF/LEF factor. **(E)** Quantification of positive cell populations from flow cytometry data. Error bars represent ±s.d. Paired T-test; ns – not significant, * <0.05, ** < 0.01 and *** <0.001. **(F)** Immunostaining of pHNES1 cells cultured in StemFlex for each of the TCF/LEF factors. Scale bars = 100 μm. **(G)** Flow cytometry of human naïve and primed cells for each TCF/LEF factor. **(H)** Quantification of gated cell populations for each TCF/LEF factor in human naïve and primed cells using flow cytometry. Error bars represent ±s.d. Paired T-test; Paired T-test; ns – not significant, * <0.05 and **** <0.0001.

First, we studied TCF/LEF expression in mouse naïve (i.e., ESC) and primed (i.e., EpiSCs) pluripotent stem cells. Immunostaining revealed homogeneous expression of Tcf7 and Tcf7l1 in both naïve and primed states (Fig. 1C), supported by flow cytometry (Fig. 1D). We observed a small albeit significant increase in the number of Tcf7-expressing cells between naïve and primed cells (Fig. 1D), whilst Tcf7l1 remained unchanged (Fig. 1E). Tcf7l2 in naïve ES cells was homogeneous, whereas in primed cells, we detected two separate populations, which is also confirmed with flow cytometry (Fig. 1D). Interestingly, immunostaining of Lef1 in naïve ES cells exhibited a clearly heterogenous pattern (including larger colonies, Fig. S1A), which becomes homogenous in primed cells (Fig. 1C and S1B). Flow cytometry faithfully captured these populations (Fig. 1D), confirming a significant increase in the number of Lef1^+^ cells in primed pluripotent conditions (Fig. 1E). Our observations align with previously reported expression of TCF/LEF genes in mouse naïve ES cells (Kelly et al., 2011; Moreira et al., 2017; Wallmen et al., 2012) and go beyond with this interesting novel observation of heterogenous Lef1 expression using immunofluorescence microscopy.

Expression of TCF/LEF factors in human naïve and primed pluripotent stem cells differ from those of mouse (Fig.1F). Immunostaining revealed low and heterogeneous presence of both TCF7 and TCF7L2 protein in naïve human pluripotent cells (HNES) cultured routinely in PXGL (**<u>P</u>**D03, **<u>X</u>**AV939, **<u>G</u>**ö6983 and **<u>L</u>**IF) on feeders (Fig. 1F), with a sizable population of TCF7L2^+^ cells also detected using flow cytometry (∼40%) (Fig. 1G). Further examination of TCF7L2 protein localisation confirmed a heterogenous profile in all cell lines examined: embryo-derived HNES1, HNES3 (Guo et al., 2016), chemically reset H9 (Guo et al., 2017) and a naïve induced pluripotent stem cell line (NIPSC3) (Bredenkamp et al., 2019)(Fig. S1C). TCF7L1 was largely absent except for a very small population (∼10%) with low expression and LEF1 was undetectable (Fig. 1G). We also assayed TCF/LEF expression in HNES cells cultured briefly under feeder-free conditions. Geltrex was either spiked in (added directly to media) at passaging or precoated on plates. HNES cells cultured on either feeders or when Geltrex was added at plating maintained their characteristic dome-shaped morphology, whereas on Geltrex-coated plates, colonies collapsed with large patches of epithelialisation (Fig. S1D). RT-qPCR of key naïve markers, such as *KLF4* and *TCFP2L1,* remain unchanged with only a small increase in *KLF17, SUSD2, DPPA3, NANOG* and *OCT4* levels when plated feeder free (Geltrex, spiked) (Fig. S1E). TCF/LEF expression remained largely unchanged with only a small increase in *TCF7* (Geltrex, spiked) and *TCF7L2* (Geltrex, coated) when compared to cells cultured on feeders (Fig. S1F). Immunostaining of naïve ES cells on feeders vs Geltrex conditions confirmed homogenous expression of TFCP2L1 with no visible changes in TCF7 and TCF7L2 expression (Fig. S1G).

Next, we investigated the expression of the TCF/LEF factors in primed human ES cells (Established H9 and capacitated HNES1 and HNES3 human naïve ES cells). When cultured routinely in commercial StemFlex medium or AFX (primed pluripotent conditions), we observed homogenous levels of nuclear-localised TCF7L1 and TCF7L2 (Fig. 1F). At a population-wide level, TCF7L1 expression was generally higher and more homogenous than that of TCF7L2 (Fig. 1 G-H), corroborating previous observations (Sierra et al., 2018). In stark contrast to mouse primed cells however, we detected very low TCF7 and LEF1 was undetectable (Fig. 1F), supported by flow cytometric analysis (Fig. 1G and H).

Taken together, these data clearly demonstrate distinct expression patterns for the TCF/LEF factors between mouse and human pluripotent stem cells, which may account for their differing requirements of WNT signalling for maintaining a naïve pluripotent state (Guo et al., 2017; Takashima et al., 2014; Ying et al., 2008). These differences between human and the mouse paradigm in the expression of TCF/LEF in naïve and primed states of pluripotency motivate and encourage a more detailed analysis in human cells of TCF/LEF expression in differentiation of extraembryonic and embryonic lineages.

### TCF/LEF expression in hypoblast cells

To study TCF/LEF factor expression in human hypoblast development, we utilised a reproducible and simple 2D differentiation protocol suitable for HNES routinely cultured in PXGL (Dattani et al., 2024, and Fig.2A). Cells were sequentially switched from 24 hours in PDA83 to 48 hours in FGF4, A8301 and XAV939. After 72 hours, patches of epithelised cells emerged surrounding small refractile colonies (Fig. 2A). Immunostaining confirmed co-expression of the hypoblast markers PDGFRA and SOX17 and not in naïve ES cells (Fig. 2B). Downregulation of naïve pluripotency markers *KLF17*, *NANOG* and *OCT4* and an upregulation of *GATA6*, *GATA4* and *SOX17*, including the hypoblast-specific markers *PDGFRA* and *APOA1,* was validated using RT-qPCR (Fig. 2C), confirming a hypoblast identity. Further qRT-PCR analysis showed a strong upregulation of *TCF7L2*, whilst *TCF7L1* and *LEF1* remained unchanged and *TCF7* was reduced in hypoblast cells compared to naïve cells (Fig. 2D). Immunostaining of hypoblast cells indicated TCF7L2 expression was low in PDGFRA^+^/SOX17^+^ cells and more prominent in a population of cells that do not express PDGFRA or SOX17 (Fig. 2E). This observation suggests a persistent population of cells expressing TCF7L2, which may represent trophectoderm, an alternative cell type also specified with this protocol (Dattani et al., 2024). Flow cytometry confirmed a small percentage of cells expressing GATA4^+^ hypoblast cells (∼28%) after 72 hours in differentiation conditions (Fig. 2F), which largely corroborated our immunostaining observations. Of note, a very small population of GATA4^+^/LEF1^+^ cells were detected accounting for 2.5-3.5% of the total population (Fig.2F).

**Fig. 2:**
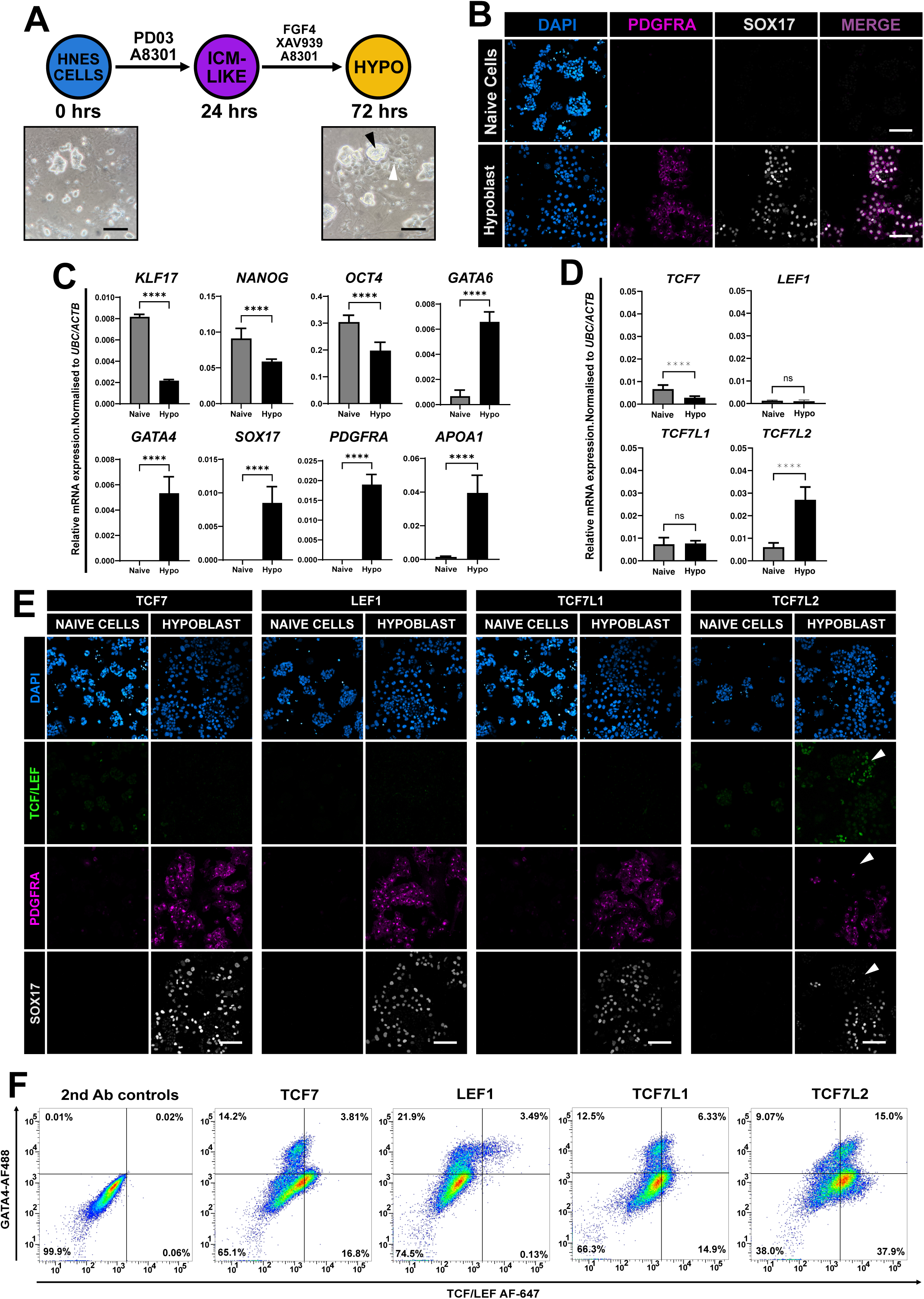
TCF/LEF expression in hypoblast cells. **(A)** Schematic of differentiation protocol to hypoblast-like cells from human naïve ES cells over 72 hours. Brightfield images of HNES3 cells 24 hours after plating (0hrs) and hypoblast-like cells in differentiation conditions 72 hours later. Undifferentiated pluripotent colony (black arrows) surrounded by nascent hypoblast-like cells (white arrows). **(B)** Immunostaining of undifferentiated HNES3 cells in PXGL and hypoblast cells after 72 hours for PDGFRA (magenta) and SOX17 (grey). Scale bar = 100 μm. **(C)** qRT-qPCR analysis of pluripotency and hypoblast markers after 96 hours either in PXGL (control) or differentiation conditions. Error bars represent ±s.d from N=3 biological replicates. Statistical analysis was performed using a paired t-test. P-values - **** = <0.001. **(D)** qRT-PCR analysis of the *TCF/LEF* genes after 96 hours either in PXGL (control) or differentiation conditions. Error bars represent ±s.d from N=3 biological replicates. Statistical analysis was performed using a paired t-test. P-values - **** = <0.001. ns – not significant. **(E)** Immunostaining of HNES3-derived hypoblast-like cells for each TCF/LEF factor (green), PDGFRA (magenta) and SOX17 (grey). Note increased TCF7L2 localisation (white arrowheads). Representative images from N=3 independent experiments. Scale bars = 100 μm. **(F)** Flow cytometric analysis of HNES3-derived hypoblast-like cells after 96 hours in differentiation conditions for each of the TCF/LEF factors. Representative of at least N=3 independent experiments.

Human naïve pluripotent cells treated with ACTIVIN A, CHIR99021 and LIF (ACL) for 5-6 days generate naïve primitive endoderm (nEND)(Linneberg-Agerholm et al., 2019) or yolk sac-like cells (Mackinlay et al., 2021), which appear to represent different stages of hypoblast development in the human embryo. Therefore, we compared TCF/LEF factor and key marker gene expression between hypoblast and nEND. When naïve ES cells were exposed to ACL for 5-6 days, naïve colonies collapsed and widespread epithelialisation was observed (Fig. S2A). We then utilised RT-qPCR to quantify mRNA transcripts between each culture condition (Fig. S2B). *KLF17, NANOG* and *OCT4* expression was all strongly reduced in nEND cells, and less strongly in hypoblast cells, corroborating previous reports (Dattani et al., 2024; Linneberg-Agerholm et al., 2019). Whilst *GATA6* and *GATA4* remained relatively unchanged between each culture condition, we observed much higher levels of *PDGFRA* and *APOA1* in hypoblast cells. *SOX17* was undetectable in nEND cells, which, interestingly, also express elevated levels of the mesoderm marker *ALPNR* (Jackson et al., 2021; Lam et al., 2023; Sagrac et al., 2020)(Fig. S2B). Further analysis revealed *TCF7L2* was clearly increased in hypoblast cultures, whereas *TCF7, LEF1* and *TCF7L1* were highly expressed in nEND, demonstrating a key difference in these cell populations (Fig. S2C). Immunostaining of hypoblast and nEND cells confirmed TCF7 in both GATA4^+^ and NANOG^+^ nuclei while LEF1 was restricted to GATA4^+^ nEND cells, and TCF7L1 was co-expressed with nests of NANOG^+^ pluripotent cells (Fig. S2D).

### Trophectoderm cells express high TCF7L2

Human naïve pluripotent cells can differentiate into trophectoderm cells in differentiation medium PDA83, containing PD03 (a MEK/ERK inhibitor) and A8301 (a potent TGFβ/NODAL inhibitor) (Guo et al., 2021; Io et al., 2021). HNES cells were plated onto Geltrex-coated plates and switched to PDA83 for 72 hours (Fig. 3A). Characteristic patches of epithelialised cells emerged and immunostaining of GATA3 expression and absence of NANOG confirmed differentiation (Fig. 3B). *KLF17, NANOG* and *OCT4* mRNA levels decreased over 72 hours in PDA83 with a concomitant strong increase in expression of the trophectoderm markers *GATA3, GATA2* and *CDX2* (Fig. 3C). When analysing TCF/LEF expression with RT-qPCR (Fig.3D), we observed reduced levels of *TCF7*, no apparent changes in *LEF1* and a strong increase in *TCF7L2* expression. Immunostaining of trophectoderm cultures at 72 hours (Fig. 3E) confirmed our initial RT-qPCR observations, including low and heterogeneous TCF7L1/GATA3^+^ cells (white arrows) and largely homogenous TCF7L2 nuclear localisation in both NANOG^+^ and GATA3^+^ cells. A small population of TCF7L1/GATA3^+^ cells (∼20%) and two populations of cells expressing TCF7L2 with GATA3 (∼53%) and without (∼16%) were present in these cultures (Fig. 3F).

**Fig. 3:**
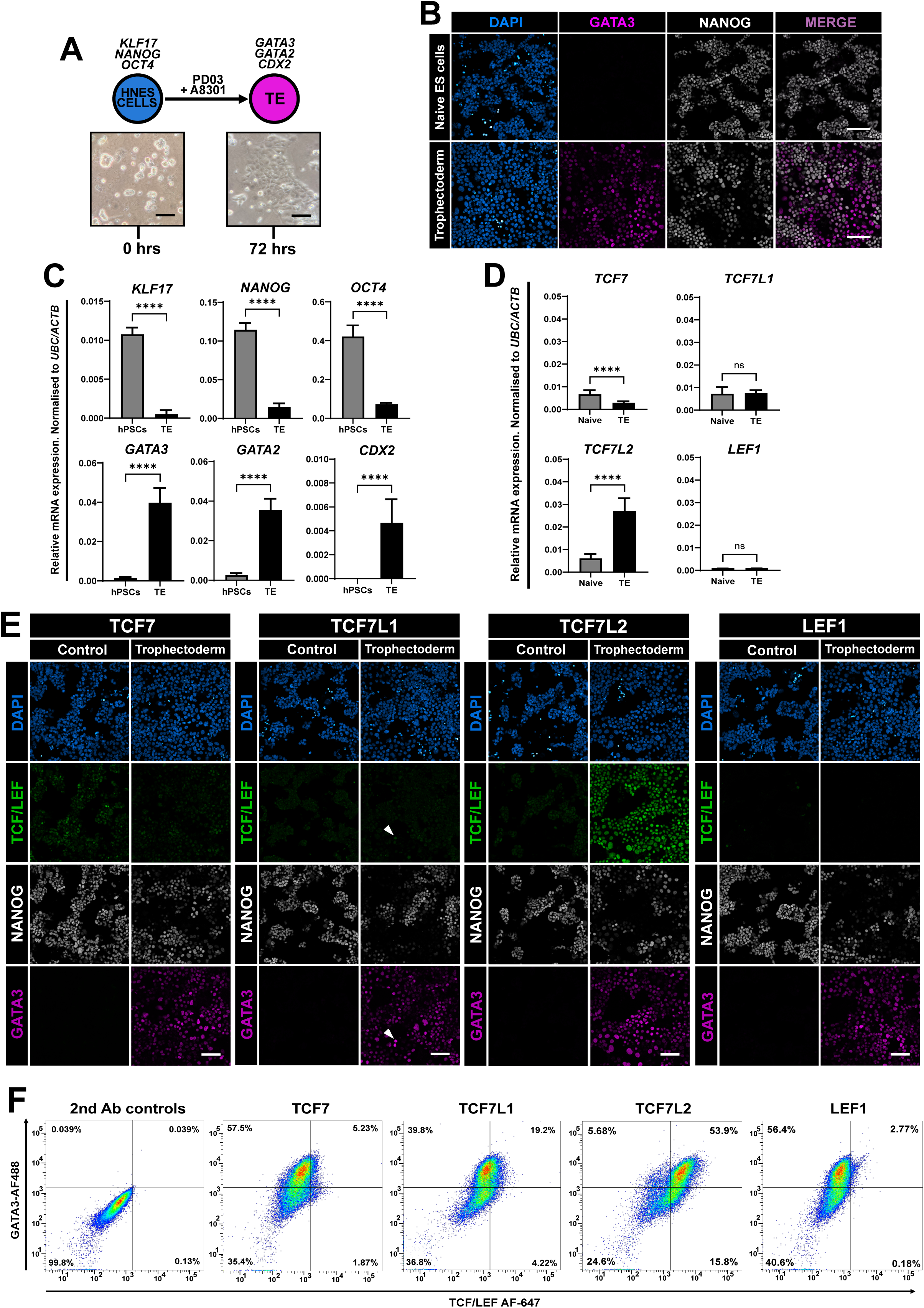
Trophoblast cells express high levels of TCF7L2. **(A)** Schematic of differentiation protocol to trophoblast-like cells from HNES cells using PD03 and A8031 for 72 hours. Brightfield images of HNES3 cells at 24 hours after plating and 72 hours later in differentiation conditions. Scale bars = 100 μm. **(B)** Immunostaining of undifferentiated HNES3 cells in PXGL and differentiated trophectoderm cells after 72 hours in PDA83 for NANOG (grey) and GATA3 (magenta). Scale bars = 100 μm. Representative of 3 fields of view and N=3 independent experiments. **(C)** qRT-PCR analysis of pluripotent and trophectoderm marker genes after 72 hours in differentiation media. Error bars represent ±s.d. Statistical analysis was performed using a paired t-test. P-values - *** = <0.01 and **** = <0.001. **(D)** qRT-PCR analysis of TCF/LEF genes after 72 hours in differentiation media. Error bars represent ±s.d. Statistical analysis was performed using a paired t-test. P-values - **** = <0.001 and ns = not significantly different. **(E)** Immunostaining of HNES3 cells cultured in PXGL and hypoblast cells after 72 hours in differentiation media for NANOG (grey), TCF/LEF factors as indicated (green) and GATA3 (magenta). White arrows represent TCF7L1 nuclei that also express GATA3. Note overlap of cells co-expressing TCF7L2 and either GATA3 or NANOG. Scale bars represent 100 μm. **(F)** Flow cytometric analysis of HNES3 and trophectoderm cells at 96 hours. Note sizeable population of cells co-expressing TCF7L2 and the trophectoderm marker (GATA3).

### WNT/β-Catenin signalling induces TCF7 and LEF1 in primed cells

WNT/β-Catenin signalling is essential for the initiation of gastrulation and subsequently the differentiation of mesendoderm cells into mesodermal and endodermal subtypes (Barrow et al., 2007; Biechele et al., 2013; Liu et al., 1999). We have established that human primed pluripotent stem cells only express TCF7L1 and TCF7L2, but neither TCF7 nor LEF1 (Fig.1D), which is in agreement with previous observations (Sierra et al., 2018) and in stark contrast to mouse primed cells (Fig.1C). We then asked whether activation of WNT/β-Catenin signalling (initiating mesendoderm differentiation) would alter TCF/LEF expression. Human primed pluripotent stem cells were treated with either CHIR or WNT3A for 24 hours. Indeed, upon CHIR or WNT3A treatment, colonies of hPSCs formed domes with refractile edges (Fig. 4A) and β-Catenin was localised to the nuclei of cells (Fig. 4B, compare inserts b1 with b2; Fig. S3A). Concomitant with a change in cell morphology, β-Catenin was present in the nuclei of CHIR-treated but not control or IWP2-treated hPSCs, similarly to previous reports in hPSCs (Dziedzicka et al., 2021; Massey et al., 2019; Sierra et al., 2018), colorectal cancer cells (Ambrosi et al., 2022) and HEK293T cells (Kafri et al., 2016). To monitor activation of the WNT/β-Catenin pathway, we measured expression of the WNT target gene *AXIN2* (Jho et al., 2002), which confirmed a strong upregulation when hPSCs were treated with CHIR and to a lesser extent with recombinant WNT3A protein (Fig. 4C and S3B). Furthermore, immunostaining established the presence of widespread and homogenous BRACHYURY protein localisation (Fig. 4D), with concomitant increased levels of *TBXT, MIXL1* and *EOMES* mRNA; strongly when treated with CHIR (Fig. 4E) and more modestly with WNT3A recombinant protein (Fig. S3B). Upon differentiation, the expression profiles of all four TCF/LEF genes changed dramatically, with both *TCF7L1* and *TCF7L2* being strongly reduced alongside an increase in *TCF7* and *LEF1* mRNA levels (Fig. 4F). There was a more pronounced effect on the expression of the TCF/LEF factors with CHIR than WNT3A, with only a small increase in *TCF7* and *LEF1* and decrease of *TCF7L1* and *TCF7L2* (Fig. S3C). Immunostaining of TCF/LEF expression in control and treated hPSCs largely corroborated our initial observations with RT-qPCR (Fig. 4G and S3D). Interestingly, we observed LEF1^+^ nuclei around the edge of the colony following WNT3A only (Fig. S3D), similarly to a previous report using micropattern gastruloids (Martyn et al., 2019). Finally, flow cytometric analysis of CHIR-treated hPSCs demonstrated a strong increase in both TCF7^+^ (∼82%) and LEF1^+^ (∼93%) cells whilst TCF7L1- and TCF7L2-expressing cell numbers were slightly decreased (Fig. 4H and 4I). Of note, despite the total number of gated cells increasing only slightly, there was an observable shift in their overall intensity (Fig. 4H).

**Fig. 4:**
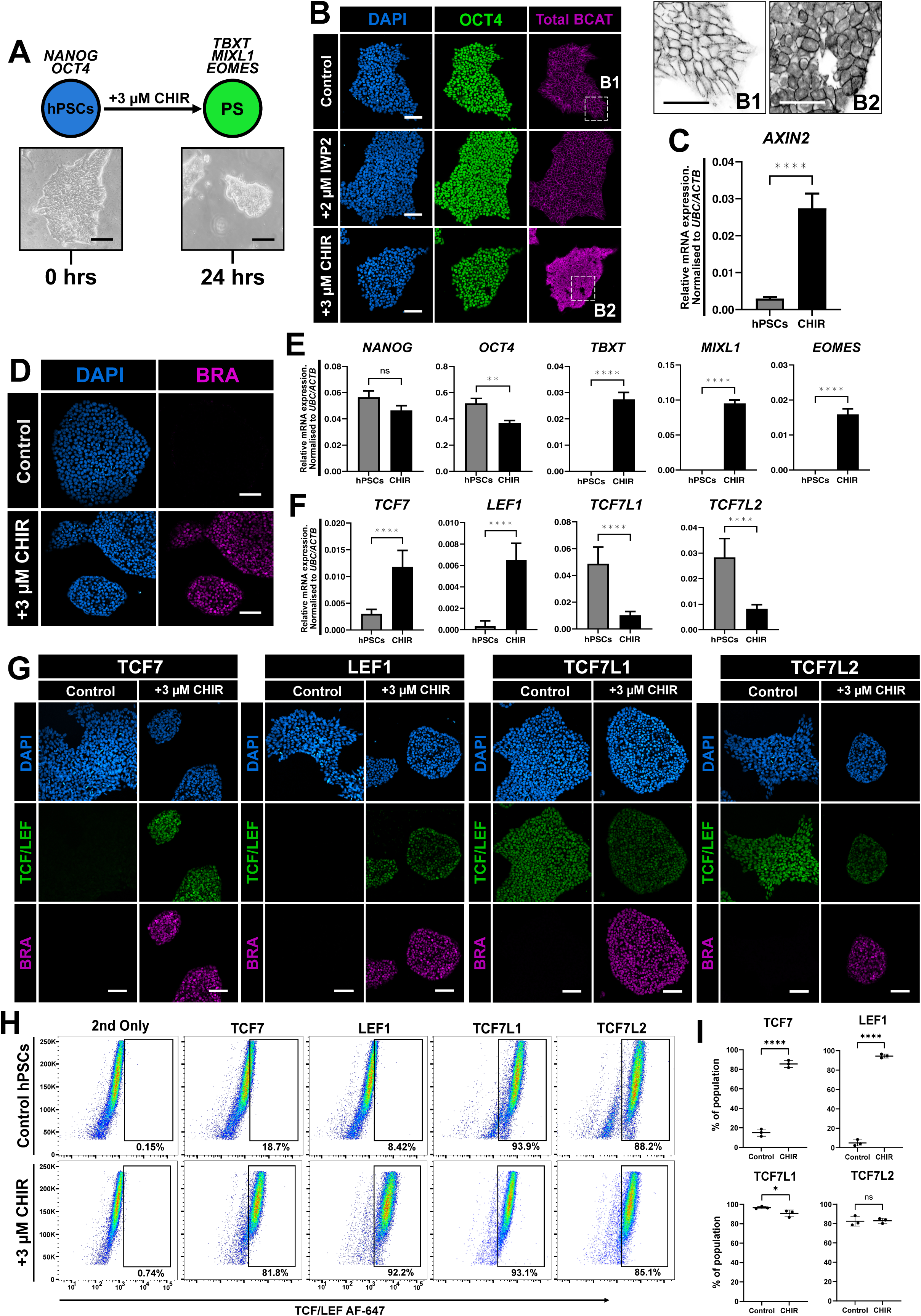
TCF7 and LEF1 are upregulated in response to WNT activation in human primed pluripotent ES cells. **(A)** Schematic of experiment of treating primed human pluripotent embryonic (ES/iPS) cells (cultured in StemFlex then switched to N2B27) with 3 µM CHIR (WNT activator) for 24 hours. Scale bars = 100 μm. **(B)** Immunostaining for β-Catenin (BCAT) (magenta) and OCT4 (green) in pHNES3 cells cultured in StemFlex medium for 72 hours following passaging with and without 2 µM IWP2 (WNT inhibitor) or switched to N2B27 + 3 µM CHIR for 24 hours. Scale bars = 100 μm. Insert images b_1_ and b_2_ are representational of each condition, respectively and scale bars = 200 um. **(C)** qRT-PCR analysis of *AXIN2* expression in control primed ES cells vs cells treated with 3 µM CHIR for 24 hours. Relative mRNA levels were normalised to *UBC and ACTB*. Student’s paired t-test (**** = *p*<0.0001). Error bars indicate ±s.d. **(D)** Immunostaining of pHNES1 cells for BRACHYURY in control and differentiation conditions (N2B27 + 3 µM CHIR for 24 hours). Scale bars = 100 μm. **(E)** qRT-PCR analysis of control vs treated primed cells for *NANOG, OCT4, TBXT, MIXL1* and *EOMES*. Relative mRNA levels were normalised to *UBC* and *ACTB*. Student’s paired t-test (ns = not significant, ** = *p*<0.01, **** = *p*<0.0001. Error bars indicate ±s.d. **(F)** qRT-PCR analysis of control vs treated primed cells for *TCF7, LEF1, TCF7L1,* and *TCF7L2*. Relative mRNA levels were normalised to *UBC* and *ACTB*. Student’s paired t-test (**** = *p*<0.0001). Error bars indicate ±s.d, N=3. **(G)** Immunostaining of pHNES3 cells treated with N2B27 + 3 µM CHIR for 24 hours for each TCF/LEF (green) and BRACHYURY (magenta). Representative images for N=5 independent experiments. Scale bars = 100 μm. **(H)** Flow cytometric analysis of control and treated (N2B27 + 3 µM CHIR for 24 hours) primed cells for each TCF/LEF factor. **(I)** Quantification of TCF/LEF expressing cells in hPSC control and differentiation conditions after 24 hours. Error bars indicate ±s.d. Student’s paired t-test (* = *p*< 0.05, *** = *p*<0.001 **** = *p*<0.0001 and ns = not significant).

Collectively, these results on TCF/LEF expression in human cells clearly differ from the mouse paradigm. In human primed pluripotent cells, we show that alongside *LEF1* (Funa et al., 2015; Martyn et al., 2019), *TCF7* is also upregulated in response to active WNT signalling. Moreover, both *TCF7L1* and *TCF7L2* are subsequently downregulated as differentiation ensues to control the spatiotemporal expression of mesendoderm genes (Hoffman et al., 2013; Sierra et al., 2018).

### TCF7L1 and TCF7L2 expression persists in amnion cells

Activation of BMP signalling in human primed pluripotent stem cells has been reported to induce their differentiation into either trophoblast (Amita et al., 2013; Roberts et al., 2014; Xu et al., 2002; Yabe et al., 2016) or amnion cells (Guo et al., 2021; Io et al., 2021; Rostovskaya et al., 2022). The embryonic relationship between these tissues remains an area of controversy but recent data suggest that primed hPSCs generate amnion, whereas naïve human embryonic stem cells give rise to trophectoderm (Guo et al., 2021; Io et al., 2021; Rostovskaya et al., 2022). We treated human primed ES cells with dual FGF and NODAL inhibition alongside BMP4 to induce amnion differentiation. After 72 hours, large sheets of flattened epithelial cells emerged (Fig. 5A), which were GATA3^+^ and NANOG^-^ (Fig. 5B). *NANOG* and *OCT4* expression levels decreased and *GATA3, GATA2, TFAP2A* and *IGFBP5* were strongly increased (Fig. 5C), denoting entry into the amnion lineage. Both *TCF7* and *TCF7L1* were decreased, *TCF7L2* was strongly increased and *LEF1* remained unchanged (Fig. 5D). Immunostaining on control and differentiated cells revealed no visible differences in TCF7, LEF1 and TCF7L1 expression, however, TCF7L2 staining appeared diminished (Fig. 5E). Flow cytometry confirmed a strong decrease in the total number of gated TCF7L2^+^ cells and less homogenous expression than in control hPSCs (Fig. 5F and G), despite an increase in mRNA levels (Fig. 5C). Delays in protein turnover compared to relative mRNA levels could account for these differences.

**Fig. 5:**
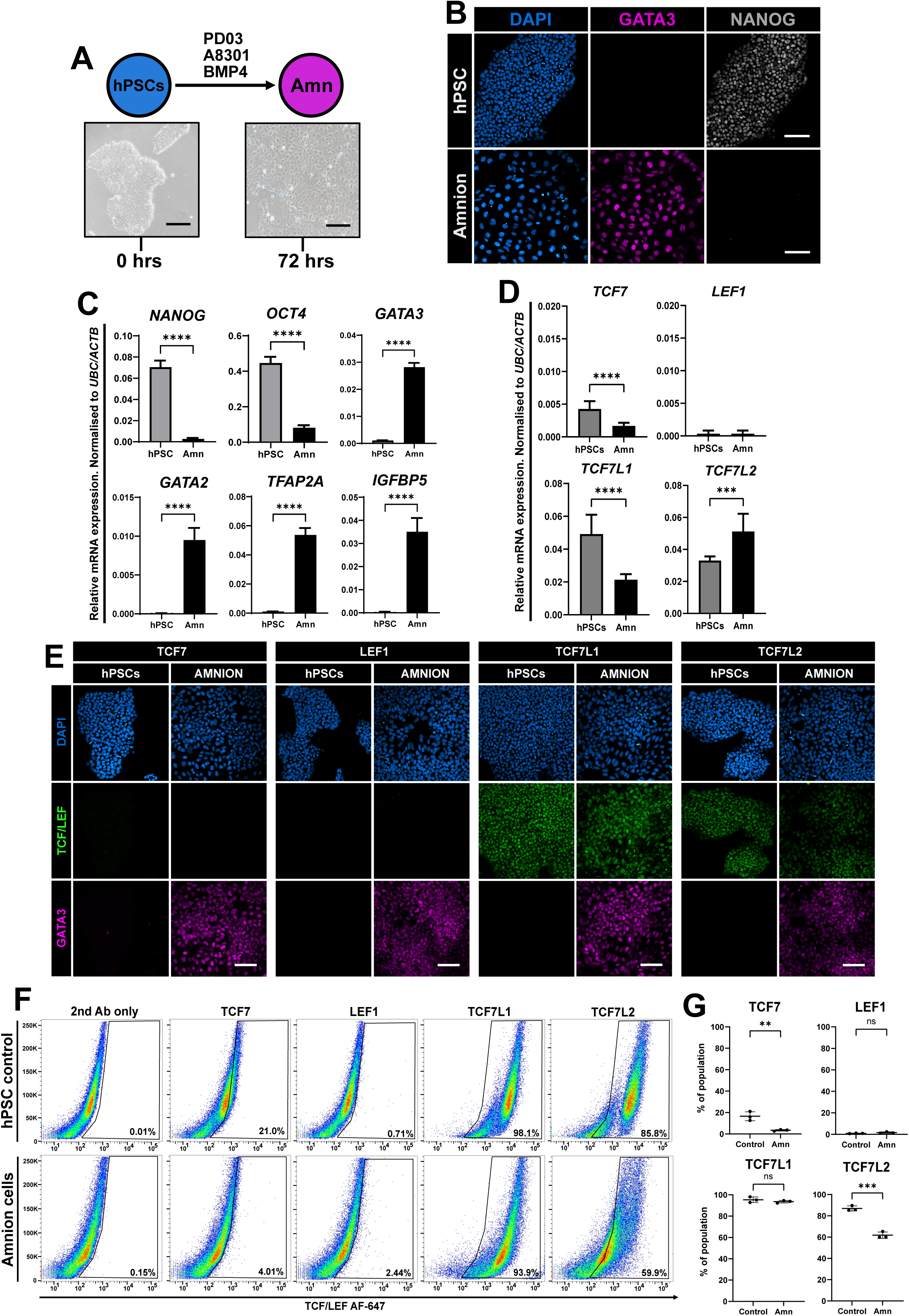
TCF7L1 and TCF7L2 remain high in Amnion differentiation. **(A)** Experimental protocol for amnion differentiation. Brightfield images of pHNES3 hPSCs and amnion cells. Scale bars = 100 μm. **(B)** Immunostaining of control pHNES3 and amnion cells after 72 hours in PDA83 + BMP4 for 72 hours of GATA3 (magenta) and NANOG (grey). Scale bars = 100 μm. **(C)** RT-qPCR of pluripotency and amnion markers from control and differentiated primed cell lines. Error bars represent ±s.d. Student’s paired t-test (**** = *p*<0.0001). **(D)** RT-qPCR of TCF/LEF factors in undifferentiated and amnion cells after 72 hours in differentiation conditions. Error bars represent ±s.d. Student’s paired t-test (*** = *p*<0.001 and **** = *p*<0.0001). **(E)** Immunostaining of control pHNES3 and differentiated amnion cells for TCF/LEF factors (green) and GATA3 (magenta). Scale bars = 100 μm. **(F)** Flow cytometry of pHNES3 and amnion cells for each of the TCF/LEF factors after 72 hours. **(G)** Gated TCF/LEF-positive cells of pHNES3 hPSCs and amnion cells quantified using flow cytometry.

These data suggest that the persistence of TCF7L1 and TCF7L2 expression may be an important prerequisite, defining an epiblast-amnion boundary for primate amnion development (Bergmann et al., 2022; Rostovskaya et al., 2022) and to shield against active WNT/β-Catenin signalling induced by BMP4 signalling (Martyn et al., 2019; Yoney et al., 2022).

### Temporal expression of the TCF/LEF factors in definitive endoderm

Shortly following the formation of the primitive streak and mesendoderm cells in the mammalian embryo (Rivera-Perez and Magnuson, 2005), fine-tuning of BMP, NODAL, WNT and FGF signalling facilitates further differentiation in mesodermal subtypes and definitive endoderm (Bardot and Hadjantonakis, 2020; Bergmann et al., 2022; Tam and Behringer, 1997; Tyser et al., 2021). We leveraged a robust protocol to efficiently differentiate primed pluripotent cells in a stepwise fashion towards definitive endoderm cells (Loh et al., 2014) and subsequently profiled the expression of the TCF/LEF factors. Human primed ES cells were treated with signalling modulators over 72 hours, which gave rise to a monolayer of epithelial cells (Fig. 6A). Whilst SOX17 was widely detectable, NANOG was lost (Fig. 6B). Pluripotency makers *NANOG* and *OCT4* were downregulated and canonical endoderm markers *SOX17, CXCR4, FGF17* and *CER1* were strongly increased at 72 hours, which was confirmed using RT-qPCR (Fig. 6C). Further analysis revealed no significant changes in *LEF1*, however, TCF7 was increased and both *TCF7L1* and *TCF7L2* were lower than in control primed ES cells (Fig. 6D). Immunostaining confirmed widespread SOX17^+^ endoderm cells with visible TCF7, TCF7L1, TCF7L2 signal and no detectable expression of LEF1 (Fig. 6E). Flow cytometric analysis revealed a population of TCF7^+^ cells (∼51%), TCF7L1^+^ decreased (∼45%) and whilst the total numbers of TCF7L2^+^ exhibited no significant difference, the overall intensity of this signal shifted (Fig. 6F). These observations were consistent over three independent cell lines in our hands (Fig. 6G, but also see recently published observations by Mukherjee et al. (2022))

**Fig. 6:**
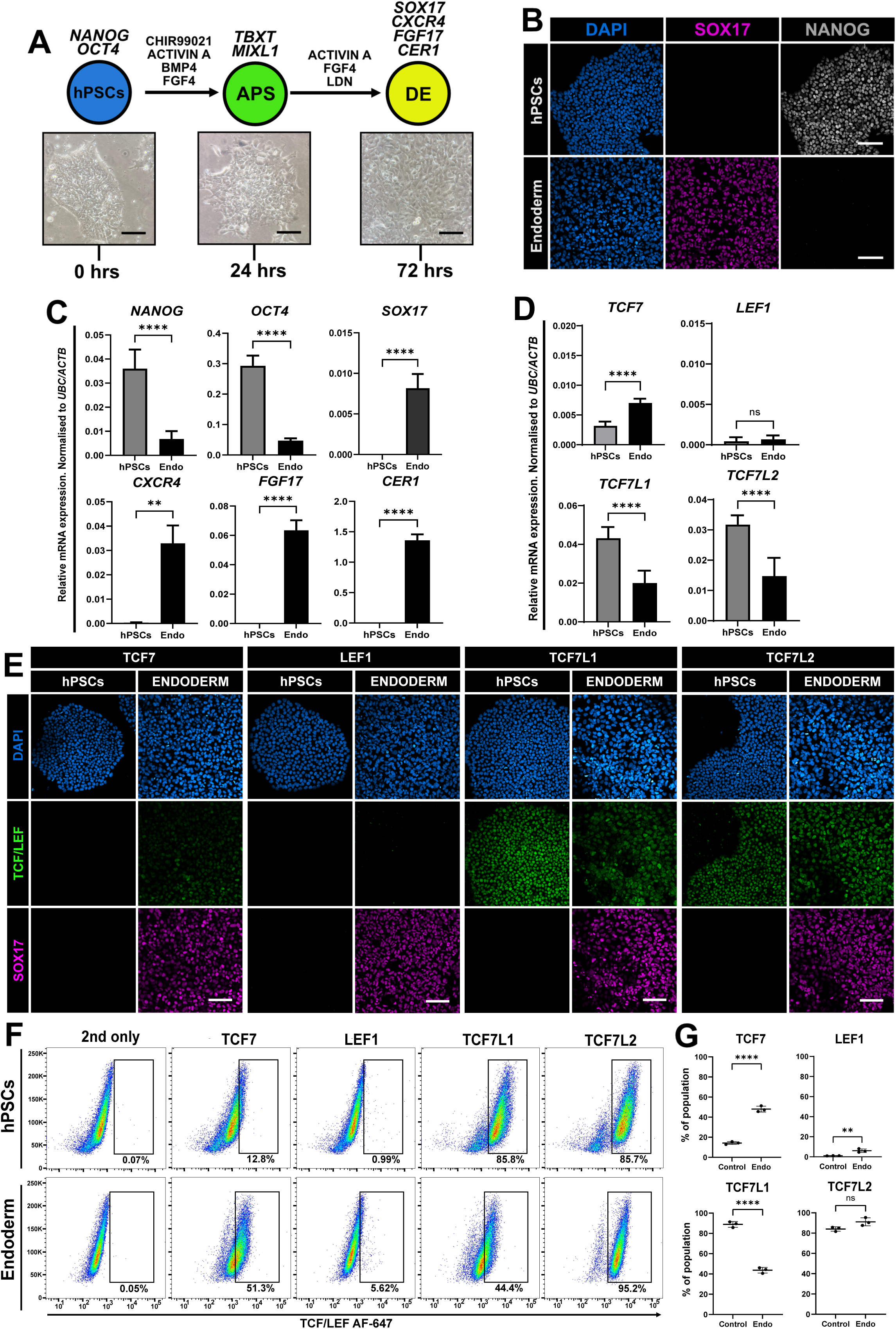
TCF/LEF factor expression in definitive endoderm from hPSCs. **(A)** Experimental schematic to differentiate human primed ES cells towards definitive endoderm. Brightfield images show control H9 and H9-derived endoderm cells. Scale bars = 100 μm. **(B)** Immunostaining of control H9 cells and H9-derived definitive endoderm after 72 hours of differentiation for SOX17 (magenta) and NANOG (grey). Scale bars = 100 μm. **(C)** RT-qPCR analysis of control vs definitive endoderm at 72 hours. Relative mRNA levels were normalised to *UBC* and *ACTB*. Student’s paired t-test (** = p< 0.01, **** = *p*<0.0001). Error bars indicate ±s.d. **(D)** RT-qPCR analysis of control vs definitive endoderm at 72 hours for *TCF7, TCF7L1, TCF7L2* and *LEF1*. Relative mRNA levels were normalised to *UBC* and *ACTB*. Student’s paired t-test (ns – not significant, **** = *p*<0.0001). Error bars indicate ±s.d, N=3. **(E)** Immunostaining of H9 hPSCs and definitive endoderm after 72 hours of differentiation for SOX17 (magenta) and each TCF/LEF factor (green). Scale bars = 100 μm. **(F)** Flow cytometric analysis of control H9 and H9-derived definitive endoderm cells for all four TCF/LEF factors after 72 hours of differentiation. **(G)** Quantification of TCF/LEF-positive cells in hPSCs and definitive endoderm samples after 72 hours. Error bars represent ±s.d. Student’s paired t-test (** =*p*< 0.01, **** = *p*<0.0001).

These findings suggest that both TCF7 and LEF1 exhibit temporal activity whereby their expression decreases over time during definitive endoderm commitment following upregulation in hPSCs after WNT stimulation. Moreover, despite the strong decreases in the mRNA levels of *TCF7L1* and *TCF7L2* in SOX17^+^ endoderm, their expression suggests a role in maintaining lineage identity, preventing precocious and differentiation into other cell types.

## Discussion

Our current understanding regarding the roles of WNT signalling underpinning early mammalian development and stem cell biology has been predominantly informed from genetic studies in rodent models. However, applying this rodent paradigm to early human development has become increasingly difficult. Here, we leveraged mouse and human pluripotent stem cell models to test conservation of expression and localisation of TCF7, LEF1, TCF7L1 and TCF7L2, the canonical terminal effectors of the WNT/β-Catenin pathway (Fig.7).

**Fig. 7:**
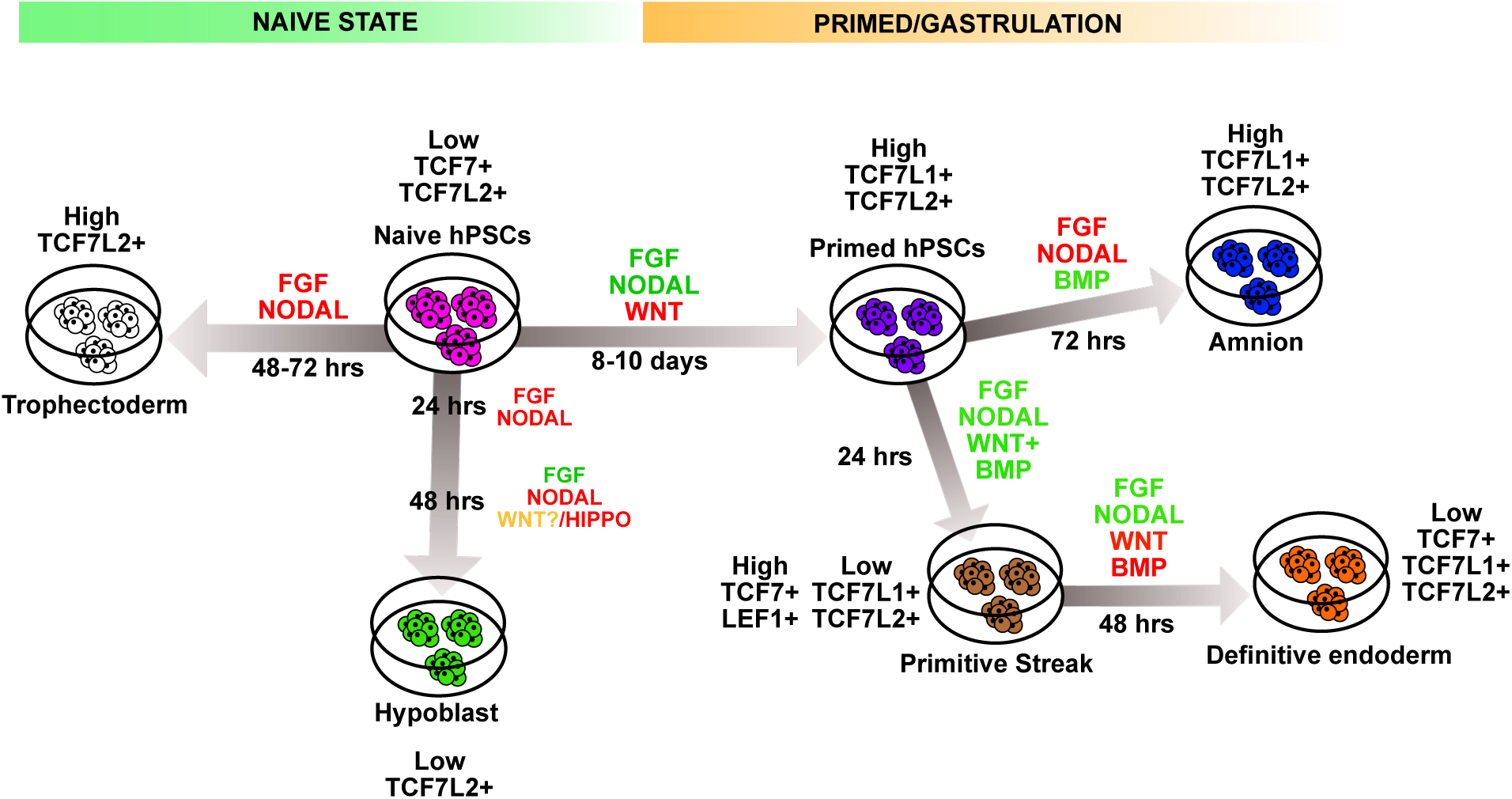
Schematic illustration of TCF/LEF factor expression in human pluripotent and differentiated cells. Summary diagram highlighting TCF/LEF transcription factor expression determined in this study using human pluripotent stem cells and lineage-specific differentiation. These experiments were conducted in cells that recapitulate cell lineages from a naïve and primed state of pluripotency. Green words = activation of signalling pathway and red words = inhibition of signalling pathway.

### Differences in states of pluripotency

We show that mouse and human comprehensively differ in expression of TCF/LEF factors in what is currently widely considered comparative states of naïve and primed pluripotency (Fig. 1). We confirm in mouse naïve ES culture the expression of all four TCF/LEF genes but find only low levels of *TCF7* and *TCF7L2* in human naïve pluripotency. It is interesting to find heterogeneous LEF1 expression in the mouse (Fig. 1C); since LEF1 expression is known to increase upon release from naïve pluripotency (Ye et al., 2017).

Rat naïve ES cells appear to express higher *Lef1* than mouse naïve ES cells and knockdown of Lef1 (Chen et al., 2013) or TALEN-mediated mutagenesis (Meek et al., 2020) can effectively counteract precocious differentiation associated with higher concentrations of CHIR. In mouse primed pluripotent stem cells (i.e., EpiSCs, Fig. 1C-E), we detect homogenous Lef1 expression, mirroring E7.5 stage embryos (Galceran et al., 2001; Oosterwegel et al., 1993). In our hands, we could not detect LEF1 in 5 different human primed ES/iPS cell lines (Fig. 1F-H, see also Fig.4F), although Mackinlay et al. (2021) have reported stronger LEF1 staining, which may be due to different culturing conditions or detection methods and likely reflects different levels of endogenous WNT signalling. Homogeneous LEF1 in mouse primed cells but not human might account for their hypersensitivity to WNT signalling. Blocking WNT signalling with either IWP2 or XAV939/IWR1 improves the derivation efficiency and long-term propagation of mouse primed cells from postimplantation embryos (Liu et al., 2017; Sugimoto et al., 2015; Sumi et al., 2013). Similarly, this approach has successfully facilitated the derivation of stable primed pluripotent stem cells from preimplantation blastocysts of pig, sheep and cow with XAV939 (Kinoshita et al., 2021b). Interestingly, combined XAV939 and IWP2 (to tightly suppress WNT signalling) was shown to derive stable rat (Iwatsuki et al., 2023) and rabbit (Kobayashi et al., 2021) primed ES cells. Finally, the inclusion of XAV939 but not CHIR captures chimera-competent ES-like cells from isolated chicken blastoderm discs (Kajihara et al., 2024). Taken together, we propose that the blockade of WNT/β-Catenin signalling in primed pluripotent cells universally serves as an important prerequisite to 1) maintain uniformity, 2) prevent premature differentiation and 3) ensure efficient directed differentiation.

These species-specific variations in TCF/LEF expression are likely related to the revealed different requirements for WNT/β-catenin signalling between mouse and human for stable propagation of the naïve pluripotent state (Bayerl et al., 2021; Bredenkamp et al., 2019; Guo et al., 2017; Lyashenko et al., 2011; Takashima et al., 2014; Wray et al., 2011; Ying et al., 2008). Culture conditions mirroring the 2i+LIF from mouse were initially found to be unsuitable for stable propagation of human naïve ES/iPS cells (Gafni et al., 2013; Hanna et al., 2010; Takashima et al., 2014). Significantly, 2i+LIF leads to strong activation of WNT signalling (via CHIR99021). Fine-tuning the concentration of CHIR drastically improves the propagation of both human (Takashima et al., 2014; Theunissen et al., 2014) and rat naïve ES cells (Chen et al., 2013; Meek et al., 2013). More recently, it has been shown that exchanging CHIR with a potent TANKYRASE inhibitor, XAV939 (initially described as a Wnt inhibitor, Huang et al., 2009), boosts self-renewal of human naïve ES cells (Bayerl et al., 2021; Bredenkamp et al., 2019; Guo et al., 2017; Khan et al., 2021; Zimmerlin et al., 2016), by predominantly buffering HIPPO signalling (Dattani et al., 2022). HNES cells lacking β-Catenin protein can self-renew and express key naïve pluripotent markers, implying that activation of the WNT/β-catenin pathway might be largely dispensable for supporting human naïve pluripotency (Bayerl et al., 2021; Dattani et al., 2022). However, we cannot rule out the possibility of compensatory mechanisms from Plakoglobin, a homologue of β-Catenin (Mahendram et al., 2013). Recently, it had been shown to boost naïve pluripotency in mouse ES cells and is expressed in the epiblast of early human embryos and naïve ES cells but not primed cells (Kohler et al., 2023).

Low *TCF7L1* expression in human naïve pluripotency raises several fundamental and exciting questions regarding its role in the regulation of pluripotency in mammals. Despite its very low expression in naïve ES cells, *TCF7L1* expression gradually increases as cells undergo capacitation (transitioning from naïve to primed pluripotency)(Rostovskaya et al., 2019) and is homogenously expressed in primed ES cells (Fig. 1F-H)(see also Sierra et al., 2018). These lower levels of TCF7L1 may contribute to the unrestricted lineage potential of human naïve ES cells (Dattani et al., 2024; Guo et al., 2021; Linneberg-Agerholm et al., 2019), even though Tcf7l1 does not restrict the differentiation of mouse naïve ES cells towards primitive endoderm (Athanasouli et al., 2023). Given its primary role as a strong repressor of gene expression (Kim et al., 2000; Liu et al., 2005; Yi et al., 2011), TCF7L1 fundamentally functions to tightly control differentiation and other signalling pathways, one of which is NODAL (Sierra et al., 2018). Human naïve and primed pluripotent cells require active NODAL signalling and its inhibition downregulates key pluripotency markers such as *NANOG* (James et al., 2005; Osnato et al., 2021; Vallier et al., 2005). When both NODAL and MEK-ERK signalling are blocked simultaneously, naïve ES cells readily undergo trophectoderm differentiation (Guo et al., 2021; Io et al., 2021). Conversely, in primed cells, TCF7L1 acts as a rheostat for a plethora of key gastrulation genes and its knockdown drastically increases *NODAL* expression (Sierra et al., 2018). Therefore, lower levels of TCF7L1 may permit high *NODAL* expression required for long-term self-renewal of human naïve ES cells. Future experiments will aim to express *TCF7L1* in HNES cells to see if they behave more like mouse naïve ES cells. Furthermore, inhibition of Nodal signalling in mouse ES cells does not impair their self-renewal in a naïve state, however, trilineage differentiation in a primed state is severely compromised (Mulas et al., 2017).

Since we have uncovered fundamental differences in TCF/LEF factor expression in human pluripotent cells between the naïve and primed pluripotent states, a future detailed analysis during capacitation is required. In support of this, the transition for naïve to primed pluripotency has revealed a fundamental requirement for WNT inhibition to circumvent precocious differentiation (Rostovskaya et al., 2019). Thus, it is of interest to understand how the epiblast transitions between pluripotent states is sufficiently shielded from WNT signalling, which may rely on SFRP family members secreted from the nascent epiblast and hypoblast (Blakeley et al., 2015; Stirparo et al., 2018). Furthermore, while we compared mouse and human in what is currently widely considered equivalent stages of pluripotency (naïve and primed), our findings here provide further evidence that they may not be equivalent in mouse and human. While both mouse and human naïve ES cells developmentally recapitulate the preimplantation epiblast of the embryo (Boroviak et al., 2014; Guo et al., 2016; Takashima et al., 2014), Mouse EpiSCs more likely represent a mid-gastrula stage (Kinoshita et al., 2021a; Kojima et al., 2014; Tesar et al., 2007; Tsakiridis et al., 2014) compared to human primed cells, which closely resemble the pre-gastrula epiblast (Tyser et al., 2021; Xiang et al., 2020). Comparative studies investigating the expression profiles of the TCF/LEF across other ES cells from a range of species will provide a wider perspective for understanding the fundamental differences in human development compared to the rodent paradigm. The expression of Tcf7l1 in mouse naïve ES cells and rapid upregulation of Lef1 present in primed cells might also be related to developmental pace. The epiblast of mouse embryos transitions from a naïve state and undergo gastrulation within 48 hours (Nichols and Smith, 2009). In humans and non-human primate embryos, the equivalent stages can take 7-10 days (Boroviak and Nichols, 2017; Rostovskaya et al., 2022; Rostovskaya et al., 2019). Thus, a lack of TCF7L1 and LEF1 expression in human naïve pluripotency may permit a longer periimplantation period required for specifying extraembryonic mesoderm and amnion, both of which emerge following gastrulation during mouse embryogenesis (Boroviak and Nichols, 2017; Ross and Boroviak, 2020).

### Extraembryonic and early embryonic cell lineages

Comparing hypoblast cells (Dattani et al., 2024) with nEND cells (naïve Endoderm, Linneberg-Agerholm et al., 2019) reveals clear differences in that hypoblast cells express only low levels of TCF7L2, whereas nEND differentiation upregulates both *TCF7* and *LEF1*. Interestingly, following the nEND protocol, we detected the upregulation of *APLNR*, usually considered a key mesoderm receptor involved in cardiac development (Jackson et al., 2021; Lam et al., 2023; Sagrac et al., 2020). Moreover, we observed a very small fraction of GATA4/LEF1^+^ cells following hypoblast differentiation (Dattani et al., 2024). Although the embryonic origin of the extraembryonic mesoderm (ExMe) remains unclear, a hypoblast origin in the peri-implantation primate embryo has recently been proposed (Ross and Boroviak, 2020). The observed low population of GATA4/LEF1^+^ cells may therefore represent ExMe differentiation from the hypoblast. Consistent with this notion, data analysis from Bergmann et al. (2022) shows high *LEF1* expression in cells demarcated to be ExMe from single cell RNA-seq data of peri/postimplantation marmoset embryos. Moreover, cells resembling ExMe have been derived from human naïve pluripotent cells (Oldak et al., 2023; Pham et al., 2022) with culture conditions containing WNT-pathway activating CHIR. Taken together, our comparative data suggest that different culture conditions induce separate populations of differentiated cells that are reminiscent of hypoblast cells, representative of early and later stages, respectively (Dattani et al., 2024; Linneberg-Agerholm et al., 2019; Mackinlay et al., 2021). These differences in culture conditions could account for the contrasting TCF/LEF expression patterns in our current observations and might be related to the status of endogenous WNT signalling (i.e., WNT activation in nEND and inhibition in hypoblast cells).

Strikingly, we observed high levels of TCF7L2 in trophectoderm cells derived directly from human naïve pluripotent stem cells. TCF7L2 has known functional links to regulation of metabolism (e.g., Verma et al., 2022), controlling context-specific cell differentiation (e.g., Lipiec et al., 2020) and proliferation (e.g., Wenzel et al., 2020). Moreover, this gene encodes alternatively spliced isoforms (reviewed by Hoppler and Waterman, 2014; and in Torres-Aguila et al., 2022) that contribute to this diverse functionality of TCF7L2 (e.g., Young et al., 2019). By controlling trophoblast cell differentiation, TCF7L2 may have ultimately a role in human embryo implantation. Intriguingly, the WNT inhibitor DKK1 is upregulated in the endometrium of women who have been hyper-stimulated with oestrogen/progesterone prior to egg collection, thereby reducing pregnancy rates (Liu et al., 2010). These increased levels of DKK1 may disrupt the embryonic-endometrial interface that is required normally to facilitate proper implantation. Our results suggest that TCF7L2 is upregulated in response to trophectoderm lineage induction, maybe prior to onset of *GATA3* expression and that heterogenous TCF7L1 expression might reflect a mixture of differentiated trophoblast subtypes (Pollheimer et al., 2006).

Positive feedback regulation of TCF7 and LEF1 following experimental WNT activation in primed human pluripotent stem cells is reminiscent of some other tissues, since these TCF/LEF genes have previously been found to be positively regulated by WNT signalling, for instance in the context of colorectal cancer (Li et al., 2006; Mayer et al., 2020; Roose et al., 1999). Both Tcf7 and Lef1 contribute to paraxial mesoderm development, somite formation and limb development (Galceran et al., 1999), with mutants phenocopying Wnt3a-null mice (Takada et al., 1994). Here, we added TCF7 to LEF1 expression induced by active WNT/β-catenin signalling using either CHIR or recombinant WNT3A protein has been described previously (Funa et al., 2015; Martyn et al., 2019). Interestingly, forced overexpression of TCF7 but not LEF1 promotes the differentiation of human primed pluripotent stem cells towards an endoderm fate by upregulating key differentiation genes such as *TBXT, MIXL1* and *GATA6* (Sun et al., 2017). Moreover, knockdown of GATA6 but not GATA4 blocks differentiation mediated by TCF7. TCF7 persistence in the endoderm (albeit at low levels) and no LEF1 in our model suggests a temporal window of induction that may also be lineage-specific and requires further in-depth examination.

Our analysis here provides a basis for future experiments to investigate upstream mechanisms of gene regulation of TCF/LEF genes during early human development, comparing it to mouse. This analysis also provides a basis for formulating novel hypotheses about regulation of downstream gene expression by WNT signalling and any possible differences between TCF7-, LEF1-, TCF7L1-, or TCF7L2-mediated downstream gene expression. How these described differences in TCF/LEF deployment between the human and mouse paradigm may account for differences in subsequent gene expression (such as for instance esrrb, Martello et al., 2012) requires further investigation. This analysis has predominantly focused on genes expression of all four TCF/LEF, however, particularly TCF7 and TCF7L2 are known to encode different mRNA and protein isoforms (Torres-Aguila et al., 2022), which may fine-tune the transcriptional response to WNT/β-Catenin signalling in early human development.

## Limitations of the study

Human embryos constitute a precious resource, requires a HFEA licence in the United Kingdom and regulatory approval prior to experimentation. Although our study was performed using pluripotent stem cells and differentiated subtypes, they only represent experimentally accessible models of early human embryogenesis. Therefore, future research will aim to perform immunofluorescence microscopy in early-stage human embryos to confirm these observations. We profiled the expression of the TCF/LEF factors in human naïve pluripotent stem cells routinely cultured in PXGL (Bredenkamp et al., 2019; Guo et al., 2017) and did not further explore alternative conditions, such as t2iLiGö (Takashima et al., 2014) or 5i/LA (Theunissen et al., 2014) that contain a low concentration of GSK3 inhibitor. CHIR is a potent inhibitor of GSK3 activity (Wu et al., 2015), an experimental tool widely used as a potent activator of downstream WNT/β-Catenin signalling. We observed more pronounced effects of CHIR (increasing BRACHYURY expression, *TCF7* and *LEF1*) than recombinant WNT3A protein when human primed cells were treated and CHIR treatment might therefore not always recapitulate a physiological response to WNT signalling. Except for *TCF7L1*, the remaining members of the TCF/LEF family exhibit extensive alternative splicing, which may be undetected by our current antibodies, accounting for differences between mRNA and protein levels. Finally, here we focussed on canonical WNT/β-catenin signalling, future research should include studying functions of non-canonical WNT signalling mechanisms in stem cell models of early human development.

## ACKNOWLEDGEMENTS

We would like to thank all members of the Hoppler and Nichols groups for critical feedback and helpful discussions on this project and Yvonne Turnbull for lab management. We thank Dr Debbie Wilkinson (Aberdeen Advanced Microscopy and Histology), Ann Wheeler (Edinburgh Imaging) and Darran Clements (Cambridge imaging) for their assistance with confocal and widefield microscopy. We thank Dr Andrea Holme (Ian Fraser Cytometry Centre, University of Aberdeen) for her kind assistance with flow cytometry.

## Funding Acquisition

This study was supported from BBSRC project grants BB/S018190/1 and BB/Y001974/1.

## AUTHOR CONTRIBUTIONS

CR and SH conceptualised this study with input from JN and TA. CR and TA performed experiments. MS designed and validated the TCF/LEF primers used for qPCR and RG provided tissue culture assistance. CR carried out all analysis and generated figures. CR and SH wrote the manuscript with input from TA, MS, RG and JN. All authors reviewed the final manuscript prior to submission.

## Materials and Methods

### Human and mouse stem cell lines

The HNES1, HNES3 (Guo et al., 2016), cR-H9 (Guo et al., 2017) and NIPSC3 (Bredenkamp et al., 2019) cell lines were a generous gift from Professor Austin Smith (Living Systems Institute, University of Exeter) after granted regulatory approval by the UK Stem Cell Committee. WA07 and WA09 were obtained from WiCell (Madison, Wisconsin). IMR-90 and KOLF2.1J human iPS cell lines were a kind gift from Dr Eunchai Kang (Institute of Medical Sciences, University of Aberdeen). E14Tg2a (Doetschman et al., 1987), wildtype AinV18 ES cells were a kind gift from Todd Evans (Turbendian et al., 2013) and the JNST wildtype cell line was derived in-house previously (Kraunsoe et al., 2023). All cell lines were all expanded in culture conditions that reflect their states of pluripotency and routinely tested for mycoplasma contamination in-house using PCR and tested negative. DNA content analysis was performed to monitor the karyotypic status of all cell lines involved in this study.

### Mouse embryonic stem cell culture

JNST #32, E14Tg2a and AinV18 (WT) mouse embryonic stem cells were cultured on 0.15% gelatin-coated plates in 2i+LIF to maintain a state of naïve pluripotency, which consisted of N2B27 supplemented with 1 µM PD0325901 (Tocris), 3 µM CHIR99021 (Tocris) and 10 ng/mL mouse recombinant LIF (Cambridge Stem Cell Institute or Peprotech). Cells were routinely cultured and passaged using Accutase (BioLegend) every 2-3 days. Epiblast stem cells (EpiSCs) were cultured to maintain primed pluripotency on 10 µg/mL bovine plasma fibronectin (Sigma Aldrich) coated plates in AFX medium which consisted of N2B27 supplemented with 10 ng/mL human/mouse/rat recombinant ACTIVIN A (Peprotech), 12 ng/mL FGF2 (Cambridge Stem Cell Institute) and 2 µM XAV939 (Tocris). EpiSCs were passaged using Accutase every 2-3 days depending on growth rate and initial seeding density. 10 µM ROCKi (Tocris) was added to N2B27-AFX medium prior at passaging for 24 hours.

### Human naïve pluripotent stem cell culture

HNES1, HNES3, cR-H9 and NIPSC3 cell lines were routinely maintained on gamma-irradiated DR4 or CF1 mouse embryonic fibroblasts (iMEFs) in N2B27 supplemented with 1 µM **<u>P</u>**D0325901, 2 µM **<u>X</u>**AV939, 2 µM **<u>G</u>**ö6983 and 10 ng/mL human recombinant **<u>L</u>**IF, termed **PXGL** as previously described (Bredenkamp et al., 2019; Guo et al., 2017). HNES cells were passaged every 3 to 4 days during exponential growth using Accutase for 12 minutes at 37°C and plated in PXGL with 10 µM Y-27635 for 24 hours. To improve attachment rate, Geltrex was diluted 1:4 with ice DMEM and 30 µL was added per 6-well. Cells used for experiments were cultured for no more than 6 consecutive times.

For feeder-free culture, stock Geltrex solution was prediluted 1:4 with ice cold DMEM and gently inverted. For coating plates, 1 mL of ice cold DMEM was added to each 6 well plate with 40 µL of prediluted Geltrex solution. Plates were incubated overnight at 4°C and then 1 hour at 37°C prior to use. Alternatively, 50 µL of diluted Geltrex solution was “spiked” into PXGL+Y at plating.

### Capacitation of human naïve ES cells

HNES1 and HNES3 cells were capacitated as previously described (Rostovskaya et al., 2019). Briefly, HNES1 and HNES3 cells were plated at 40,000 cells per 6-well in PXGL+Y for 24 hours. The next day, cells were washed once with DPBS (Gibco) and switched to N2B27 supplemented with 2 µM XAV939 (Tocris) for 10 days. After which, cells were passaged using Accutase (Biolegend) onto diluted Geltrex (1:100) coated plates in AFX or StemFlex (Thermo) medium for at least 3 passages before being used for experimentation.

### Primed/conventional human pluripotent stem cell culture

pHNES1, pHNES3, H9 embryonic stem cell lines were routinely cultured on Geltrex-coated plates (1:100 dilution overnight at 4°C then 1 hour at 37°C before use) in StemFlex medium. Confluent cells were propagated using Accutase for 5 minutes at 37°C and replated in StemFlex medium supplemented with 10 µM Y-27635 for 24 hours.

### Hypoblast differentiation

Differentiation of HNES cells towards hypoblast cells was performed as previously described (Dattani et al., 2024). HNES cells were seeded on Geltrex-coated plates in PXGL with 10 µM ROCKi for 24 hours. The next day, the cells were washed twice with 1x PBS solution and the medium was exchanged to N2B27 supplemented with PDA83 (HNES1; 1.5 µM PD03 and 1.5 µM A8301) (HNES3 and cR-H9; 1 µM PD03 and 1 µM A8301) for 24 hours. The medium was then exchanged to N2B27 supplemented with 50 ng/mL human recombinant FGF4 (Thermo), 1 µM A8301 and 1 µM XAV939 for a further 48 hours until the end of the assay. Note that the HNES1 cell line exhibits greater variability in their inductive ability to form hypoblast at the expense of trophectoderm.

### Trophectoderm differentiation

HNES cells were induced to differentiate into trophectoderm cells as previously described (Guo et al., 2021). Naïve ES cells on day 3 or 4 (depending on growth) were passaged onto Geltrex-coated plates in PXGL+Y at a seeding density of 8-10,000 cells per cm^2^. The next day, cells were washed once with PBS and switched to differentiation medium that consisted of N2B27 supplemented with PD0352901 and A8301 at titrated concentrations relevant to each cell line as follows: HNES1 (2 µM PD03 and A8301), HNES3 and cR-H9 (1.5 µM PD03 and A8301). The differentiation medium was refreshed daily and monitored for hallmark signs of differentiation (widespread epithelisation after 24 hours).

### nEnd differentiation

HNES cells were differentiated into naïve endoderm (nEND) cells as previously described (Linneberg-Agerholm., 2019; Mackinlay et al., 2021). To reduce cell death under feeder-free conditions (Geltrex-coated plates), the concentration of ACTIVIN A was reduced to 50 ng/mL. HNES cells were seeded on Geltrex-coated plates in PXGL with 10 µM for 24 hours. The next day, the cells were washed twice with 1x PBS exchanged to N2B27 supplemented with 50 ng/mL ACTIVIN A, 3 µM CHIR99021 and 10 ng/mL human LIF for 5-6 days. Medium was refreshed daily until the end of the assay.

### Immunostaining of samples

Ibidi-treated 8-well chamber slides (Ibidi) were used for immunofluorescence microscopy. Cells were fixed with 4% paraformaldehyde solution for 20 minutes at room temperature (RT) and subsequently washed three times with 1x PBS. Samples that were not processed immediately were stored at 4°C in PBS. Permeabilisation was performed using 0.5% TritonX-100-PBS for 25 minutes at RT and then blocked (5% Donkey Serum (Sigma-Aldrich), 5% BSA (Sigma-Aldrich) with 0.25% Triton-X in PBS) for 1 hour at RT on an orbital shaking platform. Primary antibodies were diluted accordingly in blocking solution and incubated overnight at 4°C. The next day, samples were washed three times with PBST (PBS with 0.02% Tween-20) and incubated with secondary antibodies/DAPI in blocking solution for 2 hours at RT on an orbital shaking platform set at 65 rpm. Samples were washed three times with 1x PBST and finally, changed to PBS prior to imaging.

### Confocal microscopy

All images were acquired using either a Leica SP5 or LSM-880 confocal microscope with either Leica, Nikon Elements or ZEN Black software, respectively. To minimise crosstalk between the channels, laser intensities were set and not adjusted throughout each imaging session. Imagining emission spectra tracks were reduced to limit bleed through. To normalise or improve the signal-noise ratio for each fluorophore, the digital gain was adjusted accordingly. TCF/LEF expression was normalised to human and mouse primed samples and TCF7/LEF1 was cross-examined with human primed ES cells treated with CHIR for 24 hours. These image settings were saved and used for each experiment and were not altered. Signal to noise reduction was performed to decrease the non-specific binding of the AF488 anti-mouse secondary antibody to unspecified components of Geltrex which was used to coat the Ibidi imaging chambers.

### Intracellular flow cytometry

Preparation of samples was performed as previously described (Festuccia and Chambers, 2011) with minor modifications. Cells were dissociated using Accutase and resuspended in ice cold 1% PFA and fixed for 1 hour at 4°C. Samples were washed once with 1x PBS and permeabilised with 70% chilled methanol-PBS solution for 1 hour on ice. Blocking was performed using 5% donkey serum, 5% BSA in PBS for 1 hour at 4°C. Primary antibodies (1:100 dilution) were added directly to blocking buffer and incubated for 1 hour. Samples were washed three times with 1:5 diluted blocking buffer. Secondary antibody staining (1:1000 dilution) was performed for 1 hour at 4°C and were subsequently washed three times. Finally, samples were resuspended into 300 μL of wash buffer and stored at 4°C for analysis (BD Fortessa X-20). Data analysis was carried out using Flow Jo software. For each experiment, a minimum of 20,000 single cells (events) were gated.

### RNA extraction, cDNA synthesis and qRT-PCR

Samples were lysed in 6-well plates using Lysis buffer and RNA was extracting using a Qiagen RNeasy column prep kit using the manufacturer’s instructions (Qiagen). RNA was eluted into RNA storage solution (Thermo) and was evaluated for integrity and signs of degradation by mixing 5 µL with 5 µL of 2x RNA buffer (Thermo) and electrophoresed for 25 minutes on a 1% agarose gel. Three distinct bands representing ribosomal subunits and tRNAs were always present with no smearing. Samples with smearing or no clear tRNA band were disposed. 500 ng of RNA was routinely reversed transcribed using SuperScript IV (Thermo) as per of the manufacturer’s instructions, however, only 0.5 µL of SSIV RT was used per reaction. cDNA was then diluted 1:10 with nuclease-free water. To ensure complete reverse transcription, 2 µL of diluted cDNA was mixed with DreamTaq 2x Master mix to amplify *GAPDH* and ran on a 1% gel for 30 minutes at 125 V. Only one distinct band was observed of equal intensity.

qRT-PCR was performed using Power Up SyBR Green MasterMix (Applied Biosystems) in 384-well white qPCR plates (Applied Biosystems) as technical quadruplicates (5 µL reactions). cT values were measured using the LightCycler II machine. cT values were obtained using the delta delta cT method and the average of *UBC* and *ACTB* were used as internal reference genes for normalisation of relative expression. Expression data were analysed using Prism Graphpad V10 along with statistical analysis. Unless specified, all experiments were performed with at least three independent cell lines (HNES1, HNES3 and cR-H9).

Primer pairs were tested to assess the melt curve, binding affinities and off-target amplification. Primers were used in RT-PCR reactions using OneTaq master mix (NEB) and PCR products were electrophoresed on a 1% agarose gel. Primers which generated a single PCR product at the predicted size were used. Several primer pairs were redesigned due to single nucleotide polymorphisms (SNPs) that interrupted annealing of the primers and reduced their efficiency. All primer pairs used here were obtained from previously published resources, the Harvard primer bank or designed using Primer-blast software. Their forward and reverse sequences are shown in table two.

### Statistics and reproducibility

All statistical analyses were carried out using GraphPad Prism V10. Individual details of statistical methods used for each experiment are presented in each figure legend. Immunofluorescence microscopy experiments were performed in at least three independent cell lines; HNES1, HNES3 (Guo et., 2016) and cR-H9 (Guo et al., 2017) and images presented were representative of 5 fields of view. qRT-PCR experiments were performed on all three independent cell lines with technical quadruplicates. Flow cytometry experiments were performed on all three independent cell lines and at least 20,000 events were recorded. In our hands, we observed comparatively lower levels of hypoblast cells and more efficient trophectoderm differentiation in the HNES1 cell line.

Antibodies denoted * in table S1 (TCF7, TCF7L1, TCF7L2 and LEF1) from Santa Cruz exhibited varying levels of activity and were difficult reproduce initial observations. Therefore, care should be taken when using these antibodies.

## Supplemental Material

**Table S1.** Antibodies used in this study.

| Target | Vendor | Item code | Dilutions |
| --- | --- | --- | --- |
| **OCT4** | Santa Cruz | SC-5279 | 1:200 (IF) |
| **NANOG** | R&D | AF1997 | 1:200 (IF) |
| **KLF17** | Atlas Antibodies | HPA024629 | 1:200 (IF) |
| **KLF4** | CST | 4038S | 1:100 (IF) |
| **TFCP2L1** | R&D | AF5726 | 1:50 (IF) |
| **GATA3** | eBioScience | 14-9966-80 | 1:200 (IF) 1:100 (FC) |
| **GATA3** | Abcam | Ab1399428 | 1:200 (IF) |
| **PDGFRA** | Abcam | Ab203491 | 1:100 (IF) |
| **BRACHYURY** | R&D | AF2085 | 1:200 (IF) |
| **GATA4** | eBioScience | 14-9980-82 | 1:100 (IF) |
| **SOX17** | R&D | AF1924 | 1:200 (IF) |
| **TCF7\*** | Santa Cruz | SC-271453 | 1:100 (IF) |
| **TCF7** | Invitrogen | MA5-14965 | 1:200 (IF) 1:100 (FC) |
| **TCF7L1\*** | Santa Cruz | SC-166411 | 1:100 (IF) |
| **TCF7L1** | PGT labs | 14519-1-AP | 1:200 (IF) 1:100 (FC) |
| **TCF7L2\*** | Santa Cruz | SC166699 | 1:100 (IF) |
| **TCF7L2** | CST | 2569S | 1:200 (IF) 1:100 (FC) |
| **LEF1\*** | Santa Cruz | SC-374412 | 1:100 (IF) |
| **LEF1** | CST | 2230S | 1:200 (IF) 1:100 (FC) |
| **β-Catenin** | Abcam | Ab16051 | 1:200 |
| **Anti-mouse 488** | Invitrogen | A-21202 | 1:1000 (IF and FC) |
| **Anti-rat 488** | Invitrogen | A-21208 | 1:1000 (IF and FC) |
| **Anti-rabbit 555** | Invitrogen | A-31572 | 1:1000 (IF and FC) |
| **Anti-rabbit 647** | Invitrogen | A-31573 | 1:1000 (IF and FC) |
| **Anti-goat 647** | Invitrogen | A-21447 | 1:1000 (IF and FC) |

**Table S2.**
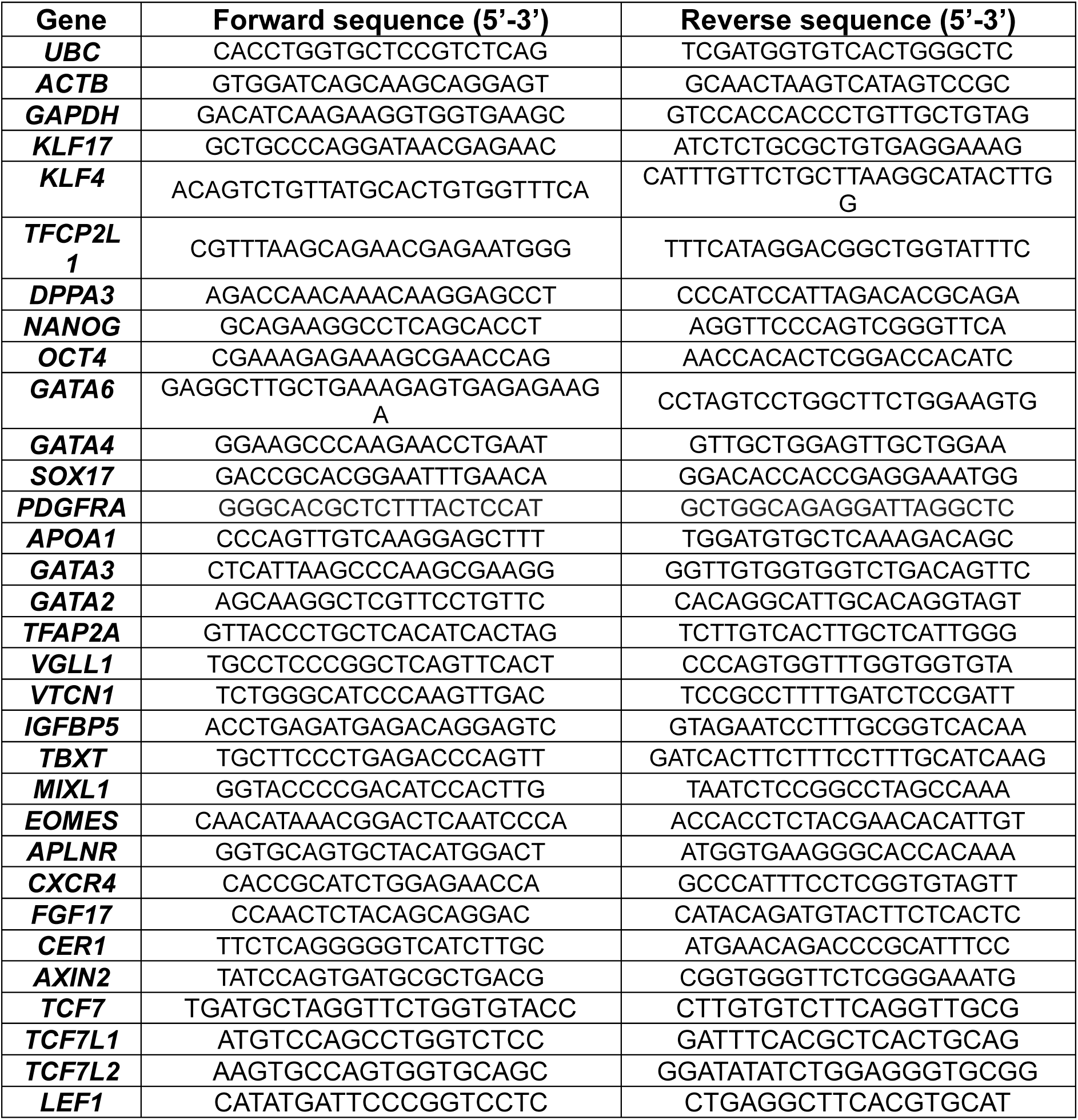
qRT-PCR primer pairs.

## Supplemental Figure legends

**Figure S1. TCF/LEF expression in mouse and human pluripotent stem cells**

**(A)** Immunostaining of JNST mouse ES cells in 2i+LIF for LEF1 (green). Scale bar = 100 μm.

**(B)** Immunostaining of JNST mouse EpiSCs cultured in AFX medium for LEF1 (magenta). Scale bar = 100 μm.

**(C)** Immunostaining of HNES1, HNES3, cR-H9 and NIPSC3 cells cultured in PXGL on feeders for TCF7L2 (green) and OCT4 (magenta). Scale bars =100 μm.

**(D)** Brightfield images of HNES3 cells cultured on feeders, spiked in Geltrex and coated wells. Black arrows represent areas of collapsed colonies and epithelialisation. Scale bars = 100 μm.

**(E)** qRT-qPCR assay of HNES1, HNES3 and cR-H9 cells cultured on feeders or with Geltrex (spiked in and coated) at 96 hours for key pluripotency markers. Relative mRNA levels were normalised to *UBC* and *ACTB*. Student’s paired t-test (* = *p<*0.05, ** = *p*< 0.01, *** = *p*<0.001 **** = *p*<0.0001 and ns = not significant). Error bars indicate ±s.d.

**(F)** qRT-qPCR of HNES1, HNES3 and cR-H9 cells cultured on feeders or with Geltrex (spiked in and coated) at 96 hours for all four TCF/LEF genes. Relative mRNA levels were normalised to *UBC* and *ACTB*. Student’s paired t-test (** = *p*< 0.01, *** = *p*<0.001 **** = *p*<0.0001 and ns = not significant). Error bars indicate ±s.d.

**(G)** Immunostaining of HNES3 cells cultured on feeders in PXGL for TCF7 or TCF7L2 (green) with the naïve markers, TCFP2L1 (magenta). Scale bars = 100 μm.

**Figure S2. Comparison of TCF/LEF factor expression in nEND and hypoblast cells**

**(A)** Experimental schematic for the differentiation of HNES cells using ACTIVIN, CHIR and LIF (ACL) for 5 days to generate nEND cells. Scale bars = 100 μm.

**(B)** qRT-PCR of naïve, nEND and hypoblast cells for pluripotency and hypoblast markers. Relative mRNA levels were normalised to *UBC* and *ACTB*. Student’s paired t-test (** = *p*< 0.01, *** = *p*<0.001 **** = *p*<0.0001 and ns = not significant). Error bars indicate ±s.d.

**(C)** qRT-PCR of naïve, nEND and hypoblast cells for all four TCF/LEF genes. Relative mRNA levels were normalised to *UBC* and *ACTB*. Student’s paired t-test (* = *p<*0.05, **** = *p*<0.0001 and ns = not significant). Error bars indicate ±s.d.

**(D)** Immunostaining of D3 HNES3-derived hypoblast cells for PDGFRA (cyan), TCF/LEF factors (green) and SOX17 (magenta) and D5 HNES3-derived nEND cells for GATA4 (cyan), TCF/LEF factors (green) and NANOG (magenta). Scale bars = 100 μm.

**Figure S3. TCF7 and LEF1 are upregulated in response to WNT activation with CHIR or recombinant WNT protein**

**(A)** Immunostaining of control or treated H9 cells (with 300 ng/mL mouse recombinant WNT3A protein or 3 µM CHIR99021) for 24 hours. Images are represented as inverted LUTs. Scale bars = 100 μm. Inserts representative for each culture condition (A1: control, A2: WNT3A and A3: CHIR). Scale bars = 200 µm.

**(B)** qRT-PCR of pluripotency and gastrulation markers after 24 hours in control and treated hPSCs. Relative mRNA levels were normalised to *UBC* and *ACTB*. Student’s paired t-test (* = *p<*0.05, ** = *p*< 0.01, *** = *p*<0.001, **** = *p*<0.0001 and ns = not significant). Error bars indicate ±s.d.

**(C)** qRT-PCR of all four TCF/LEF transcription factor genes in control hPSCs vs treated for 24 hours. Relative mRNA levels were normalised to *UBC* and *ACTB*. Student’s paired t-test (* = *p<*0.05, ** = *p*< 0.01, *** = *p*<0.001). Error bars indicate ±s.d.

**(D)** Immunostaining of H9 cells treated with either recombinant WNT3A protein or CHIR for 24 hours for TCF/LEF (grey), OCT4 (green) and BRACHYURY (magenta). Scale bars = 100 μm.

