## Supplementary figures and images for "WNT-mediating TCF/LEF transcription factor gene expression in early human pluripotency and cell lineages differs from the rodent paradigm"

### Suppl Figures

#### Supplementary Figure 1.

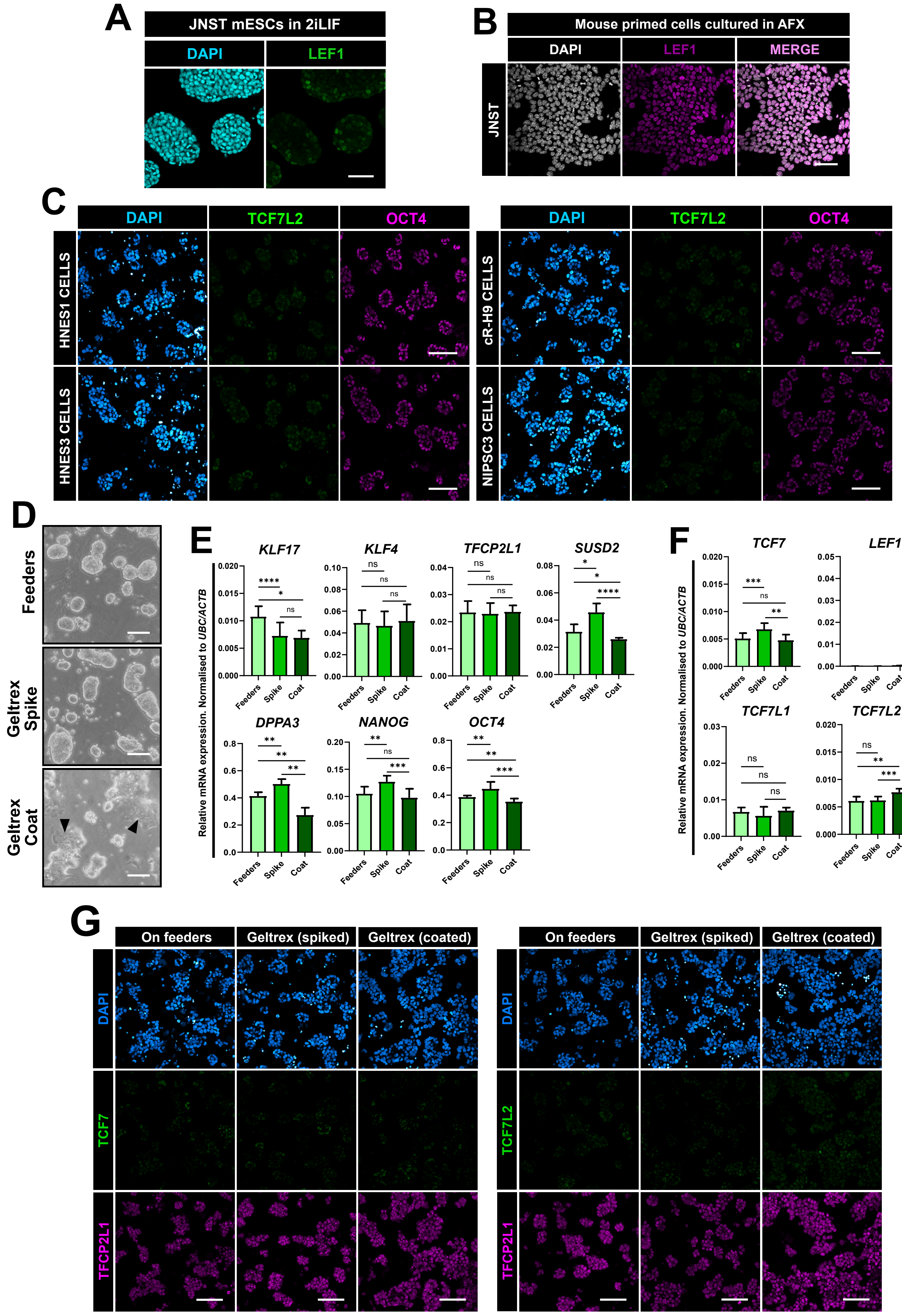

##### Supplementary figure 2.

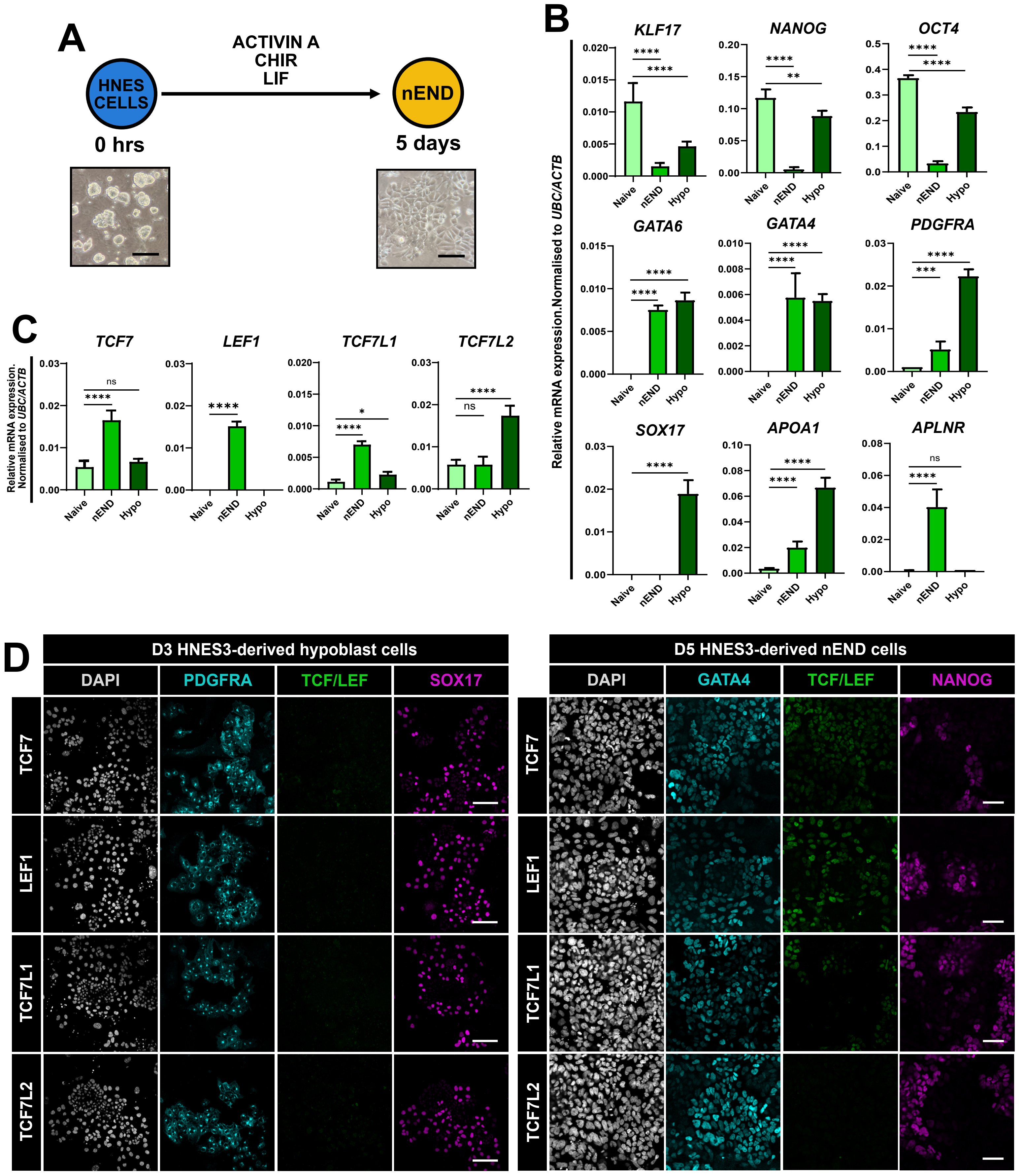

Supplementary Figure 3.

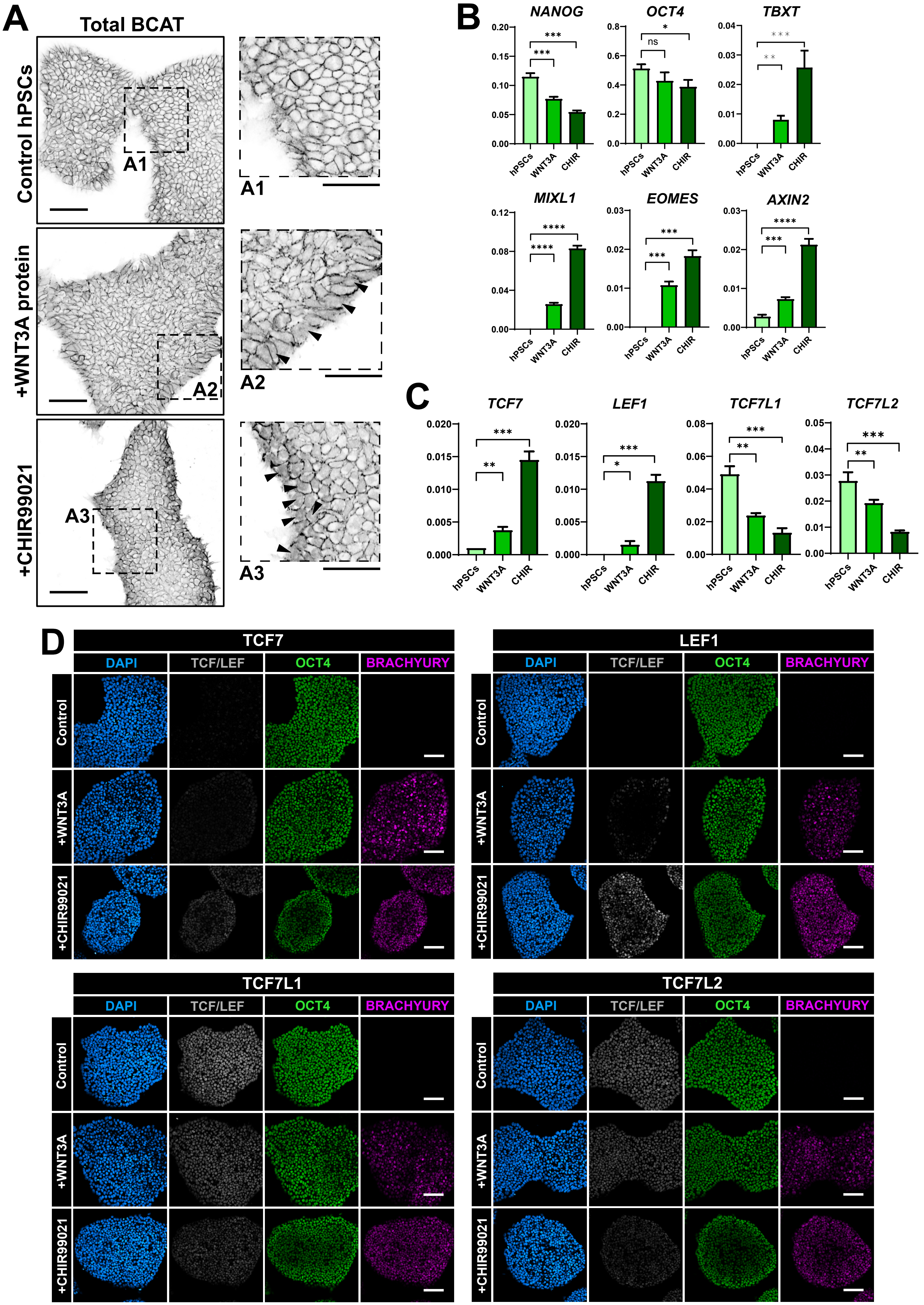
